# Bile Acids Serve As Endogenous Antagonists Of The Leukemia Inhibitory Factor (LIF) Receptor in Oncogenesis

**DOI:** 10.1101/2023.09.30.560269

**Authors:** Cristina Di Giorgio, Elva Morretta, Antonio Lupia, Rachele Bellini, Carmen Massa, Ginevra Urbani, Martina Bordoni, Silvia Marchianò, Pasquale Rapacciuolo, Claudia Finamore, Valentina Sepe, Maria Chiara Monti, Federica Moraca, Nicola Natalizi, Luigina Graziosi, Eleonora Distrutti, Michele Biagioli, Bruno Catalanotti, Annibale Donini, Angela Zampella, Stefano Fiorucci

## Abstract

The leukemia inhibitory factor (LIF) is member of IL-6 family of cytokines involved immune regulation, morphogenesis and oncogenesis. In cancer tissues, LIF binds a heterodimeric receptor (LIFR), formed by a LIFRβ subunit and glycoprotein(gp)130, promoting epithelial mesenchymal transition and cell growth. Bile acids are cholesterol metabolites generated at the interface of host metabolism and the intestinal microbiota. Here we demonstrated that bile acids serve as endogenous antagonist to LIFR in oncogenesis. The tissue characterization of bile acids content in non-cancer and cancer biopsy pairs from gastric adenocarcinomas (GC) demonstrated that bile acids accumulate within cancer tissues, with glyco-deoxycholic acid (GDCA) functioning as negative regulator of LIFR expression. In patient-derived organoids (hPDOs) from GC patients, GDCA reverses LIF-induced stemness and proliferation. In summary, we have identified the secondary bile acids as the first endogenous antagonist to LIFR supporting a development of bile acid-based therapies in LIF-mediated oncogenesis.

## Introduction

Bile acids are atypical steroids generated in the human body by the coordinate activity of liver and bacterial enzymes ^1^. In the liver, two major metabolic pathways, known as the classical and the alternative, transform cholesterol into primary bile acids, cholic acid and chenodeoxycholic acid (CA and CDCA) that, after conjugation with taurine (T) or glycine (G), are secreted in the biliary tree and then released in the intestine^3^. In the small intestine, bile salts are first deconjugated and then dehydroxylated by the intestinal microbiota, giving rise to secondary bile acids, lithocholic acid and deoxycholic acid (LCA and DCA) and ursodeoxycholic acid (UDCA)^6^. The majority of primary bile acids (95%) are reabsorbed in the terminal ileum and transported back to the liver through the entero-hepatic circulation, while the majority of LCA is excreted with the faeces^3^. Other bile acid species, that are generated at the host/microbial interface, are the 3-, 7- and 12-oxo derivatives, allo-derivatives and iso-allo derivatives^8^. The biological functions of these bile acids are still poorly understood although there is growing interest for their potential as immune/metabolic mediators^10^.

Bile acids have been linked to cancer development^12,14^. Because their amphipathic structure, high luminal content and detergent effects on cell membranes, bile acids have been historically considered as potential cancer-promoting agents in entero-hepatic tissues.^12,16^ However, the epithelial-damaging effects, underlying these pro-oncogenic properties, manifest at concentrations of bile acids > 100 µM while, at physiological intracellular concentrations (that are in the nano-micromolamolar range), bile acids are not cytotoxic and function as signalling molecules activating a family of cell membrane and nuclear receptors, known as bile acid-activated receptors, that maintains tissue and immune homeostasis. The Farnesoid-X-receptor (FXR), a receptor for primary bile acids^18^, and the G protein-coupled bile acid receptor 1 (GPBAR1)^20^, a receptor for secondary bile acids, are the two main bile acid sensors, functioning as integrative hubs between the intestinal microbiota and host metabolism and immunity^22^. Of relevance, FXR and GPBAR1 activation by natural and synthetic agonists^24,26^ confers protection against colorectal cancers development, while their genetic ablation promotes entero-hepatic tumorigenesis in animal models^28,30,31^. Together these data suggest that, while high intraluminal concentrations of bile acids in the gastro-intestinal tract promote epithelial damage and inflammation-driven metaplasia, physiological levels contribute to maintenance of epithelial homeostasis by regulating epithelial barrier integrity, maintaining intestinal stemness and immune and microbial homeostasis^32,33^. However, there is no information of bile acids intratumor concentrations in the large majority of cancers.

In addition to FXR and GPBAR1, bile acids serve as non-exclusive ligands for other nuclear receptors, including the pregnane-X-receptor (PXR)^34^, vitamin-D-receptor (VDR)^35^, peroxisome-proliferator activated receptors (PPARs)^36^, liver-X-receptors (LXRs)^37^ and the retinoid orphan-related receptor (ROR) γT^38^, and membrane receptors such as the Sphingosine 1 receptor (SP1R)2^39^ and M2/M3 muscarinic receptors ^40,41^. In these settings, bile acids function as receptor agonists and, up to now, there is no evidence that bile acids might function as direct antagonists to any receptor.

The leukaemia inhibitory factor (LIF) is a member of interleukin (IL)-6 cytokine’s family secreted by epithelial cells and monocytes^42^. LIF is a pleiotropic cytokine regulating differentiation, proliferation and survival in embryo and adult cells. LIF is also involved in cancer growth and invasiveness driving epithelial mesenchymal transition (EMT) process in pancreatic colon and gastric adenocarcinomas^5,43–46^ but also in extraintestinal cancers^47^. In target cells, LIF binds to an heterodimeric complex formed by two subunits, the LIF receptor (LIFR) and the glycoprotein (gp) 130^46^. In addition to LIF, the LIFR/gp130 heterodimer is also activated, although in a non-exclusive manner, by oncostatin M (OSM) which, in addition, transduces its signalling through a specific receptor complex made up by OSMR/gp130^48^. LIFR may also participate to the formation of tripartite receptor complexes with gp130 and the CNTF receptor activated by cardiotrophin 1 (CT-1), cardiotrophin-like cytokine factor 1, ciliary neurotrophic factor (CNTF) and neuropoietin (NP)^49,50^. Binding of LIF to the LIFR/gp130 complex leads to a gp130-dependent phosphorylation of the signal transducer and activator of transcription STAT3 that in turn is central in regulating the immune response and results activated in the majority of human cancers ^51^. In previous studies, we have reported that BAR502 ^52–54^, a semisynthetic steroidal agonist of FXR and GPBAR1, exerts a tumour suppressor effect by acting as LIFR antagonist and STAT3 indirect inhibitor ^5^.

Building on this background, we have investigated whether natural bile acids might function as LIFR antagonists. Our results demonstrate that secondary bile acids are natural LIFR antagonists identifying DCA, GDCA, TDCA and 3-oxoDCA as the most potent endogenous antagonists of LIFR. Additionally, analysis of bile acids species in GC paired samples demonstrated that the tissue expression of LIFR is inversely correlated with the tumour content of GDCA and that tissue content of GDCA declines in the GC compared with non-neoplastic paired samples. Together, these studies prove that bile acids are LIFR antagonists and that a reduced content of GDCA in GC tissues contributes to enhance LIF-LIFR-STAT3 signalling.

## Material and Methods

### Bile Acids synthesis

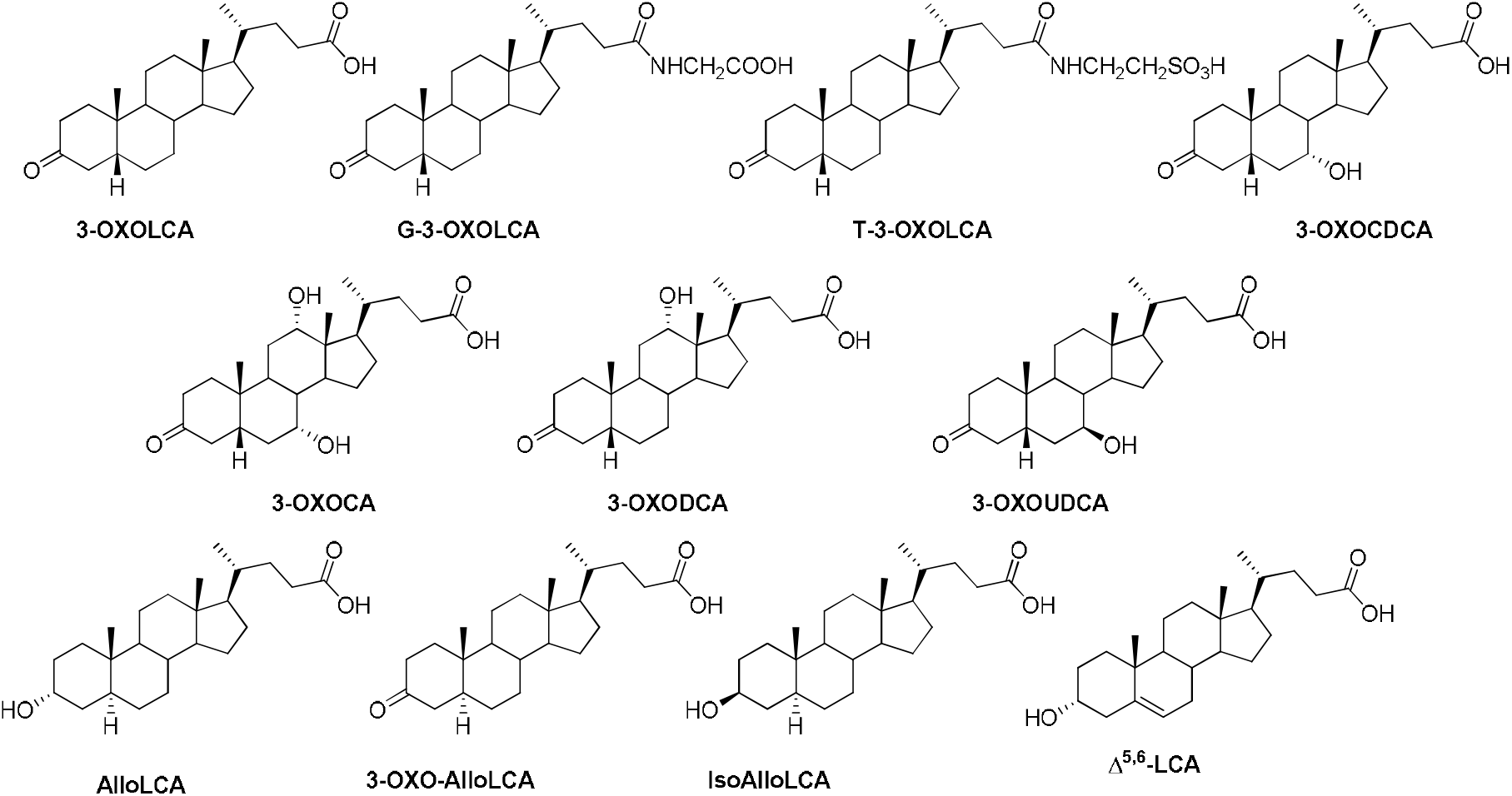

### Bile acids derivatives synthesized

Jones’ reagent was used for the regioselective oxidation of secondary alcohols, 3α,7α-dihydroxy-5β-cholanoic acid (CDCA) and 3α-hydroxy-5β-cholanoic acid (LCA), in ketones, 7α-hydroxy-3-keto-5β-cholanic acid and 3-keto-5β-cholanic acid, respectively. The reaction was conducted at 0°C with excellent yield (Scheme 1). Tauro-and glyco-conjugated 3-oxolithocholic acid were synthesized, using EDC, DIPEA and a catalytic amount of HOBt as coupling reagents for compound **T-3-oxoLCA** (quantitative yield) and subsequent basic hydrolysis to obtain with quantitative yield the compound **G-3-oxoLCA**.

**Scheme 1^a^:**
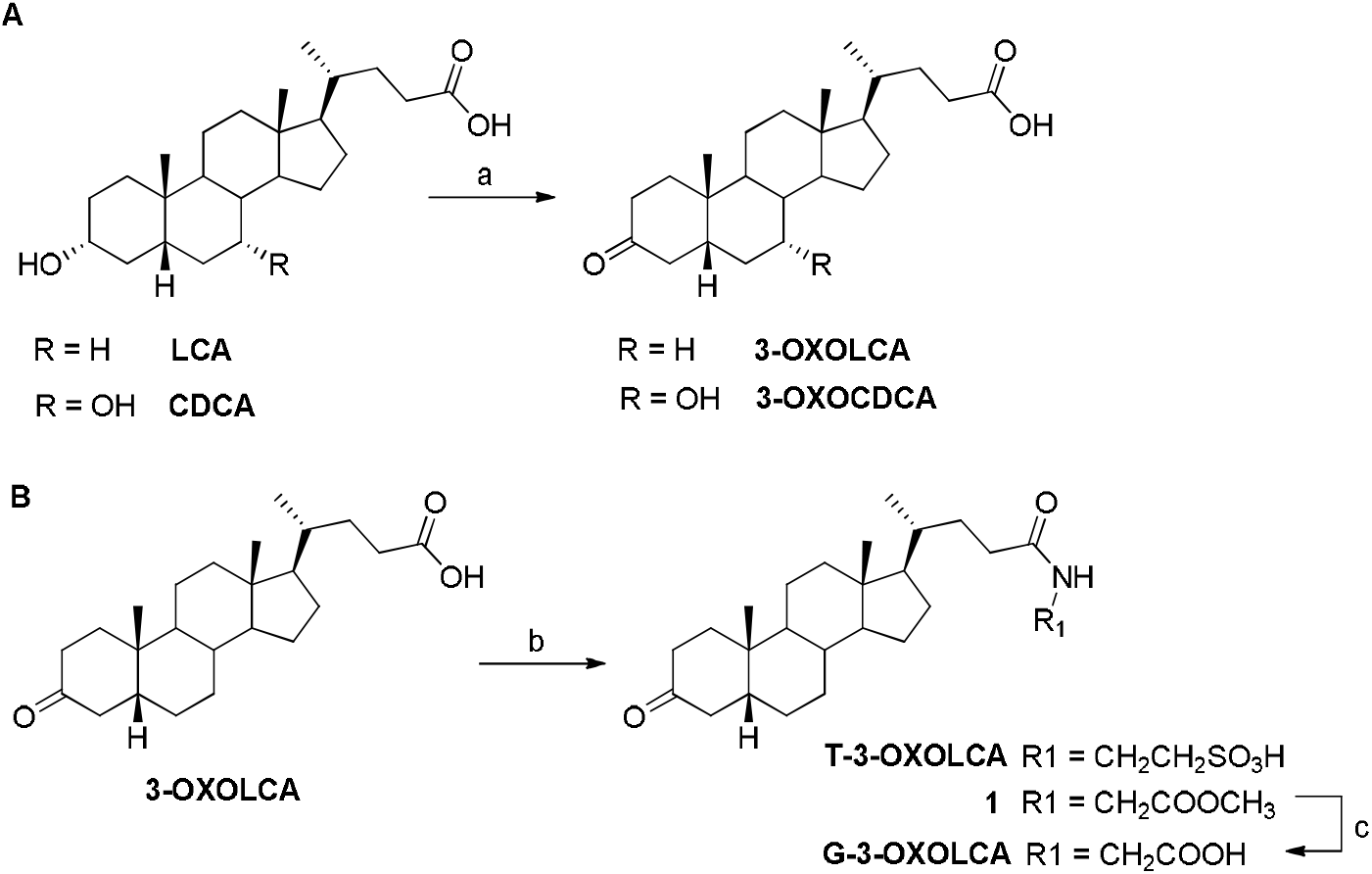
^a^Reagents and conditions: a) CrO_3_, H_2_SO_4_, Acetone, 0°C, 80%; b) EDC. HCl, HOBt, DIPEA dry, DMF dry, taurine or glycine methylester, quantitative yield; c) NaOH, H_2_O:MeOH 1:1 v/v, quantitative yield.

3-oxocholic acid (3-oxoCA) was synthesized by regioselective C3 oxidation of cholic acid using Fetizon’s reagent (scheme 2, panel A). Firstly, CA was esterified with Fisher esterification (*p*-TSA in CH_3_OH). The corresponding methyl ester was then refluxed in freshly distilled toluene with 2 equiv. of Fetizon’s reagent (silver carbonate on Celite). Crude mixture was finally submitted to alkaline hydrolysis with NaOH in CH_3_OH, affording the desired 3-ketocholic acid (90% yield).

**Scheme 2^a^:**
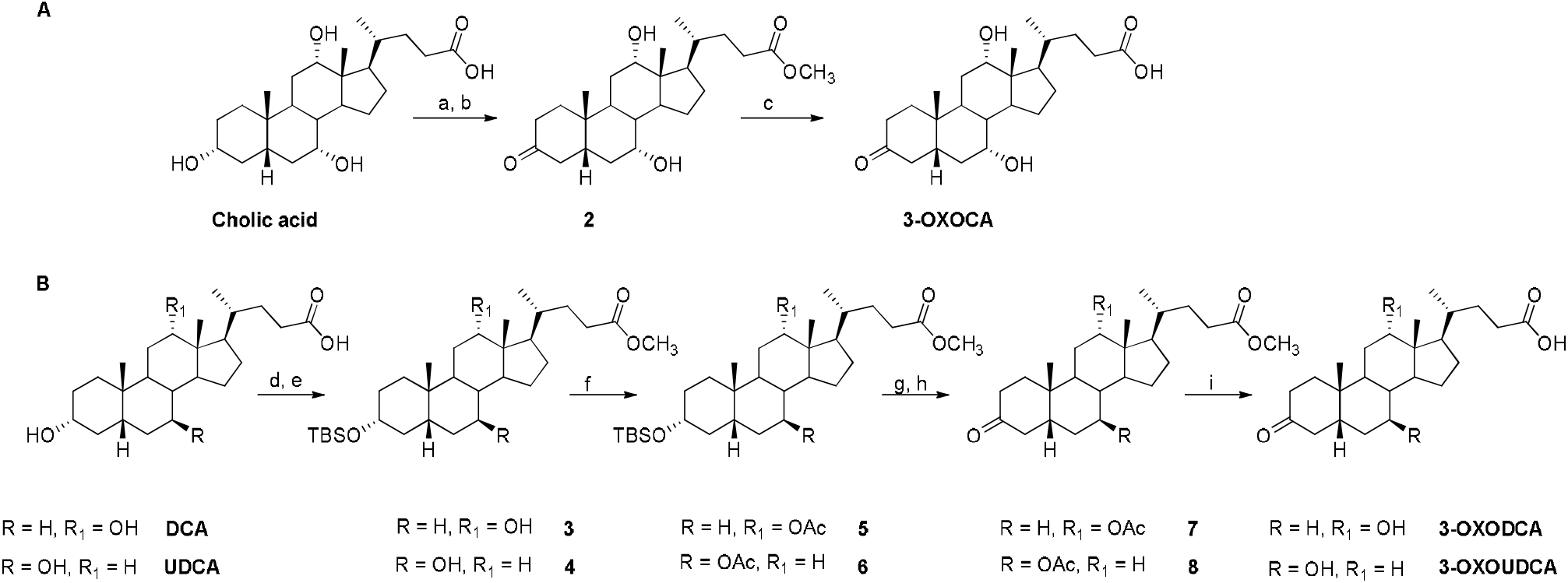
^a^Reagents and conditions: a) p-TsOH, in MeOH dry, quantitative yield; b) Ag_2_CO_3_ on celite, toluene dry, reflux, 60%; c) NaOH, MeOH:H_2_O 1:1 v/v, reflux, 90%; d) p-TsOH, MeOH dry, quantitative yield; e)

TBSCl, imidazole, in DMF dry: pyridine dry 2:1 v/v, 0°C, 60%; f) acetic anhydride in pyridine dry, 90%; g) TBAF, THF dry, 78%; h) CrO_3_, H_2_SO_4_, acetone, 70%; i) NaOH, MeOH:H_2_O 1:1 v/v, reflux, 90%.

Unfortunately, using the same oxidative condition to the deoxycholic acid (DCA) and ursodeoxycholic acid (UDCA), the reactions were not successful and very low yields were obtained. Alternatively, after TBS protection of the hydroxyl at C3, the hydroxyl at C12 and C7 of DCA and UDCA were acetylated, respectively, in order to obtain compounds **5** and **6**. After TBAF deprotection, Jones oxidation at C3, followed by alkaline hydrolysis, furnished **3-oxoDCA** and **3-oxoUDCA**.

Finally, we therefore chemically synthesized unique secondary bile acids, including Δ_5,6_-Lithocholic acid (LCA), alloLCA, 3-oxo-alloLCA, and isoalloLCA. All these secondary bile acids have a “flat” shape that results in an A/B-trans orientation. To achieve these 5α-cholane derivatives, HDCA methyl ester was firstly monoprotected at C3 with TBSCl and then activated at C6 with p-toluensolfonyl chloride, obtaining compound **9** that was subjected to elimination at C6 and then deprotection at C3 with TBAF. Hydrolysis at C24 furnished Δ**_5,6_-LCA**, while an aliquot of compound **11** was hydrogenated to afford the required A/B *trans* ring junction. Finally, hydrolysis at methyl ester gave **AlloLCA**, that was subjected to Jones oxidation in the same experimental condition previously described, to afford **3-oxo-AlloLCA**.

To obtain isoalloLCA, compound **12** was prepared in a multi-step procedure, involving ditosylation at C3 and C6, simultaneous inversion at the C3 position and elimination at the C6 position and deacetylation at C3, as previously described ^2,4^ Hydrogenation of double bond with H_2_ and Pd(OH)_2_/C and hydrolysis of methyl ester furnished **IsoAlloLCA**.

**Scheme 3^a^:**
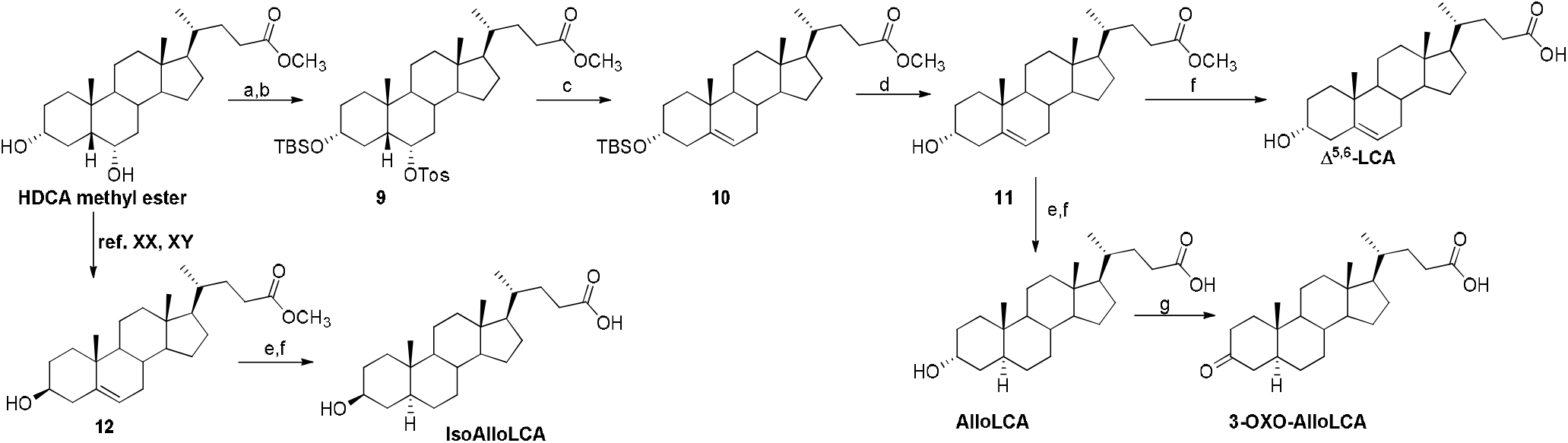
^a^Reagents and conditions: a) TBSCl, imidazole, DMF:pyridine 2:1, 0°C, 60%; b) p-toluenesulfonyl chloride in pyridine dry, quantitative yield; c) LiBr, Li_2_CO_3_ in DMF dry, quantitative yield; d) TBAF, in THF dry, quantitative yield; e) H_2_, Pd(OH)_2_/C degussa type, THF: MeOH dry 1:1 v/v, quantitative yield; f) NaOH, MeOH:H_2_O 1:1 v/v, reflux, 60-80%; g) CrO_3_, H_2_SO_4_, acetone, 0°C, 80%.

All chemicals were obtained from Sigma-Aldrich and solvents and reagents were used as supplied from commercial sources with some exception. Tetrahydrofuran and toluene were distilled from calcium hydride immediately prior to use. All reaction were carried using flame-dried glassware. Reaction progress was monitored via thin-layer chromatography (TLC) on Alugram silica gel G/UV254 plates. The purification of synthetized compounds was carried out by flash chromatography on Biotage® Selekt. NMR spectra were obtained on Bruker 400 spectrometer and recorded in CDCl_3_ (δ_H_ = 7.26, δ_C_ = 77.0 ppm). Coupling constants (*J*) are reported in hertz (Hz) and chemical shifts (δ) are in ppm and referred to CHCl_3_ as internal standards. Spin multiplicities are given as s (singlet), d (doublet), t (triplet), or m (multiplet). High-resolution ESI-MS spectra were performed with LTQ-XL equipped with an Ultimate 3000 HPLC system (Thermo Fisher Scientific) mass spectrometer.

#### Synthesis of 3-oxo-5β-cholanoic acid (**3-oxoLCA**) and 7α-hydroxy-3-oxo-5β-cholanoic acid (**3-oxoCDCA**)

To a solution of LCA (1.0 g, 2.66 mmol) and, alternatively, CDCA (1.0 g, 2.55 mmol) in acetone, 0.54 mL/mmol of Jones reagent solution (CrO_3_, H_2_SO_4_) was added at 0°. After 1 h, the mixtures were extracted with NH_4_Cl saturated solution and dichloromethane (3 × 50 mL). The combined organic layers were washed with brine, dried on Na_2_SO_4_, filtered, and evaporated to give **3-oxoLCA** and **3-oxoCDCA** (800 mg of each, 80% yield) as crude products.

#### 3-oxo-5β-cholanoic acid (**3-oxoLCA**)

An analytic sample was obtained by HPLC on a Nucleodur 100-5 C18 (5 μm; 4.6 mm i.d. x 250 mm), with MeOH/H_2_O (84:16) and trifluoroacetic acid 0.1% as eluent (flow rate 3 mL/min) (t_R_ = 12.0 min).

Selected ^1^H NMR (CDCl_3_, 400 MHz): δ_H_ 2.69 (1H, t, *J* = 14.2 Hz), 2.38 (2H, m), 2.27 (1H, m), 2.16 (1H, m), 1.02 (3H, s), 0.93 (3H, d, *J* = 6.4 Hz), 0.69 (3H, s).

^13^C NMR (CDCl_3_, 100 MHz): δ_C_ 213.7, 179.6, 56.4, 55.9, 44.3, 42.8, 42.3, 40.7, 40.0, 37.2, 37.0, 35.5, 35.3, 34.8, 30.9, 30.8, 28.1, 26.6, 25.8, 24.1, 22.6, 21.2, 18.3, 12.1. HR ESIMS m/z 373.2743 [M-H]^-^, C_24_H_37_O_3_ requires 373.2748.

#### 7α-hydroxy-3-oxo-5β-cholanoic acid (**3-oxoCDCA**)

An analytic sample was obtained by HPLC on a Nucleodur 100-5 C18 (5 μm; 4.6 mm i.d. x 250 mm), with MeOH/H_2_O (80:20) and trifluoroacetic acid 0.1% as eluent (flow rate 3 mL/min) (t_R_ = 24.4 min).

Selected ^1^H NMR (CDCl_3_, 400 MHz): δ_H_ 3.94 (1H, m), 3.40 (1H, t, *J* = 14.2 Hz), 2.41 (2H, m), 2.26 (2H, m), 1.02 (3H, s), 0.96 (3H, d, *J* = 6.4 Hz), 0.71 (3H, s).

^13^C NMR (CDCl_3_, 100 MHz): δ_C_ 213.6, 178.7, 68.6, 55.8, 50.3, 45.6, 43.2, 42.8, 39.5, 39.3, 36.9, 36.8, 35.3 (2C), 33.8, 33.3, 30.7, 30.6, 28.2, 23.7, 21.9, 20.9, 18.2, 11.8.

HR ESIMS m/z 389.2692 [M-H]^-^, C_24_H_37_O_4_ requires 389.2697.

#### Synthesis of tauro-3-oxo-5β-cholanoic acid (**T-3-oxoLCA**) and glyco-3-oxo-5β-cholanoic acid (**G-3-oxoLCA**)

To a two different solutions of 3-oxoLCA (400 mg, 1.1 mmol) in N,N-dimethylformamide anhydrous, EDC. HCl (4 eq., 4.4 mmol), HOBT (4 eq., 4.4 mmol) and N,N-diisopropylethylamine (8 eq., 8.8 mmol) were added. After 30 minutes, taurine (2 eq., 2.2 mmol) or glycine methyl ester (2 eq., 2.2 mmol) were added, and the reactions were stirred overnight at room temperature. Then the mixtures were extracted with water and ethyl acetate (3 × 50 mL). The combined organic layers were washed with an aqueous solution of KOH 2.5 M and brine, dried on Na_2_SO_4_, filtered and evaporated to give **T-3-oxoLCA** (500 mg) and a glycoconjugate methyl ester intermediate (600 mg, quantitative yield), as crude products.

The glycoconjugate methyl ester (600 mg) was subjected to alkaline hydrolysis with NaOH in H_2_O: MeOH 1:1 at reflux for 1 h, to obtain **G-3-oxoLCA** (quantitative yield) as crude product.

#### Tauro-3-oxo-5β-cholanoic acid (**T-3-oxoLCA**)

Crude **T-3-oxoLCA** (500 mg) was purified in a reverse-phase flash chromatography with octadecyl-functionalized silica gel and methanol (25 mL), as mobile phase (310 mg, 0.643 mmol, 58% yield). An analytic sample was obtained through HPLC on a Nucleodur 100-5 C18 (5 μm; 4.6 mm i.d. x 250 mm), with MeOH/H_2_O (85:15) and trifluoroacetic acid 0.1% as eluent (flow rate 3 mL/min) (t_R_ = 11.4 min).

Selected ^1^H NMR (CD_3_OD, 400 MHz): δ_H_ 3.61 (2H, t, *J* = 6.7 Hz), 2.98 (2H, t, *J* = 6.7 Hz), 2.27 (2H, m), 2.13 (2H, m), 0.97 (3H, s), 0.96 (3H, d, ovl), 0.70 (3H, s).

^13^C NMR (CD_3_OD, 100 MHz): δ_C_ 212.0, 174.2, 56.7, 56.2, 51.5, 44.3, 42.8, 42.3, 40.7, 40.0, 37.2, 37.0, 36.1, 35.5, 35.3, 34.8, 30.9, 30.8, 28.1, 26.6, 25.8, 24.1, 22.6, 21.2, 18.3, 12.1. HR ESIMS m/z 480.2785 [M-H]^-^, C_26_H_42_NO_5_S requires 480.2789.

#### Glyco-3-oxo-5β-cholanoic acid (**G-3-oxoLCA**)

An analytic sample was obtained by HPLC on a Nucleodur 100-5 C18 (5 μm; 4.6 mm i.d. x 250 mm), with MeOH/H_2_O (85:15) and trifluoroacetic acid 0.1% as eluent (flow rate 3 mL/min) (t_R_ = 24 min).

Selected ^1^H NMR (CDCl_3_, 400 MHz): δ_H_ 4.10 (2H, t, *J* = 5.2 Hz), 2.69 (1H, t, *J* = 13.8 Hz), 2.34 (2H, m), 2.18 (2H, m), 1.03 (3H, s), 0.95 (3H, d, *J* = 6.6 Hz), 0.70 (3H, s).

^13^C NMR (CDCl_3_, 100 MHz): δ_C_ 211.6, 174.0, 173.8, 56.5, 56.0, 45.3, 43.8, 43.3, 41.3, 40.5, 40.2, 37.2, 37.1, 35.5, 35.3, 34.8, 30.9, 30.8, 28.1, 26.6, 25.8, 24.1, 22.6, 21.2, 18.3, 12.1. HR ESIMS m/z 430.2961 [M-H]^-^, C_26_H_40_NO_4_ requires 430.2963.

#### Synthesis of 3-oxo-7α,12α-dihydroxy-5β-cholanoic acid (**3-oxoCA**)

Cholic acid (1.0 g, 2.45 mmol) was dissolved in dry methanol and a catalytic amount of p-toluenesulfonic acid was added. The mixture was stirred at room temperature overnight, then poured into water, neutralized with sodium bicarbonate saturated solution and evaporated under vacuum. The residue was extracted with water and EtoAc (3 x 50 mL), then the organic layer was dried on Na_2_SO_4_, filtered and evaporated to furnish a crude residue in quantitative yield (1.1 g, 2.6 mmol) that was subjected to next oxidation step without further purification.

To a solution of CA methyl ester (1.1 g, 2.6 mmol) in 100 mL toluene, Ag_2_CO_3_ on celite (2 eq.) was added in portions. The suspension of Ag_2_CO_3_ on celite was pre-refluxed for 0.5 h to be activated and completely dehydrated. The suspension was refluxed, and the water was removed by a Dean-Stark apparatus. When reaction was completed, the mixture was filtered to remove the solid, and the filtrate was concentrated in vacuum to provide a yellowish solid as crude residue. Purification was done by flash chromatography with eluent CHCl_3_: MeOH (98:2) to furnish pure C3 oxidized derivative **2** (651 mg, 1.55 mmol).

Compound **2** (651 mg, 1.55 mmol) was subjected to alkaline hydrolysis for treatment with NaOH pellets (10 eq.) in a solution of MeOH: H_2_O 1:1 v/v (20 mL) at reflux. When the reaction was completed, the resulting solution was concentrated under pressure, diluted with water, acidified with HCl 6N and extracted with EtoAc (3 x 50 mL). The collected organic phases were washed with brine, dried over Na_2_SO_4_ anhydrous, filtered, and evaporated under reduced pressure to furnish carboxylic acid 3-oxoCA as crude product (565 mg, 1.39 mmol, 90% yield), that was subjected to further purification using HPLC apparatus.

#### 3-oxo-7α,12α-dihydroxy-5β-cholanoic acid (**3-OXOCA**)

An analytic sample was obtained by HPLC on a Nucleodur 100-5 C18 (5 μm; 4.6 mm i.d. x 250 mm), with MeOH/H_2_O (80:20) and trifluoroacetic acid 0.1% as eluent (flow rate 3 mL/min) (t_R_ = 13.0 min).

Selected ^1^H NMR (CDCl_3_, 400 MHz): δ_H_ 4.05 (1H, br s), 3.94 (1H, s), 3.39 (1H, t, *J* = 13.8 Hz), 2.41 (3H, m), 2.21 (3H, m), 1.00 (3H, ovl), 0.99 (3H, ovl), 0.74 (3H, s).

^13^C NMR (CDCl_3_, 100 MHz): δ_C_ 213.7, 178.2, 72.9, 68.4, 47.0, 46.6, 45.5, 42.9, 41.8, 39.5, 36.6, 35.2, 34.9, 33.7, 31.0, 30.8, 30.7, 28.5, 27.5, 27.2, 23.2, 21.6, 17.3, 12.5.

HR ESIMS m/z 405.2643 [M-H]^-^, C_24_H_37_O_5_ requires 405.2646.

#### Synthesis of 3-oxo-12α-hydroxy-5β-cholanoic acid (**3-oxoDCA**) and 3-oxo-7β-hydroxy-5β-cholanoic acid (**3-oxoUDCA**)

**DCA** (2.0 g, 5.09 mmol) or **UDCA** (2.0 g, 5.09 mmol) were dissolved in dry methanol and a catalytic amount of *p*-toluenesulfonic acid was added. The mixtures were stirred at room temperature overnight, then poured into water, neutralized with sodium bicarbonate saturated solution, and evaporated under vacuum. The residues were diluted with water and extracted with ethyl acetate (3 x 50 mL). The combined organic layers were dried on Na_2_SO_4_, filtered, and evaporated to furnish crude residues (5.16 mmol) that were subjected to next synthetic step without further purifications.

Crude **DCA** and **UDCA methyl esters** (5.16 mmol) were dissolved in N,N-dimethylformamide:pyridine anhydrous (2:1) and then imidazole and tert-butyldimethylsilyl chloride were added at 0°C. The mixtures were stirred for 6 h at 0°C, then evaporated in vacuum and extracted for three times with water and ethyl acetate (3 x 50 mL). The combined organic layers were dried on Na_2_SO_4_, filtered, and evaporated to furnish crudes that were purified on flash chromatography with 95:5 dichloromethane/methanol, furnishing 1.55 g of pure C3 protected intermediates (compounds **3** and **4**) (3.1 mmol, 60% yield).

Derivatives **3** and **4** (3.1 mmol), were dissolved in anhydrous pyridine (3 eq.) and acetic anhydride (1 eq.) was added. The reactions were stirred at room temperature for 5 hours and then evaporated under vacuum. The crudes were extracted with water and EtOAc (3 x 50 mL), dried on Na_2_SO_4_, filtered, and evaporated to furnish compounds **5** and **6,** that were subjected to next synthetic step without further purifications.

Crude derivatives were solubilized in dry THF and then an excess of tetra-*n*-butylammonium fluoride (TBAF) solution (1M in THF dry) was added. When the reactions were completed, the residues were extracted for three times with H_2_O and EtOAc (3 x 50 mL). Then the organic layers were dried, filtered and concentrated under vacuum to give deprotected intermediates as crude residues (980 mg, 2.18 mmol, 78% yield) that were subjected to C3 oxidation without further purifications. To a solution of crude deprotected intermediates (980 mg, 2.18 mmol, 78% yield) in acetone, 0.54 mL/mmol of Jones reagent solution (CrO_3_, H_2_SO_4_,) was added at 0°. After 1h, the mixtures were extracted with NH_4_Cl saturated solution and dichloromethane (3 × 50 mL). The combined organic layers were washed with brine, dried on Na_2_SO_4_, filtered and evaporated to give derivatives **7** and **8** (1.53 mmol, 70% yield) as crude products.

Finally, derivatives **7** and **8** (1.53 mmol) were treated with NaOH pellets (10 eq.) in a solution of MeOH: H_2_O 1:1 v/v (20 mL) at reflux, to obtain the simultaneous remotion of C7/C12 acetyl group and hydrolysis of C24 methyl ester functionality. When the reactions were completed, the resulting solutions were concentrated under vacuum, diluted with water, acidified with HCl 6N and extracted with EtoAc (3 x 50 mL). The collected organic phases were washed with brine, dried over Na_2_SO_4_ anhydrous and evaporated under reduced pressure to furnish carboxylic acids **3-oxoDCA** and **3-oxoUDCA** as crude residues (1.38 mmol) in 90% yield, that were subjected to purification through HPLC.

#### 3-oxo-12α-hydroxy-5β-cholanoic acid (**3-oxoDCA**)

An analytic sample was obtained by HPLC on a Nucleodur 100-5 C18 (5 μm; 4.6 mm i.d. x 250 mm), with MeOH/H_2_O (65:35) and trifluoroacetic acid 0.1% as eluent (flow rate 3 mL/min) (t_R_ = 16.3 min). Selected ^1^H NMR (CDCl_3_, 400 MHz): δ_H_ 4.08 (1H, t, *J* = 3.0 Hz), 2.73 (1H, t, *J* = 14.2 Hz), 2.41 (2H, m), 2.28 (1H, m), 2.17 (1H, m), 1.00 (3H, s), 0.99 (3H, d, *J* = 6.4 Hz), 0.73 (3H, s). ^13^C NMR (CDCl_3_, 100 MHz): δ_C_ 214.7, 179.5, 73.4, 48.1, 47.3, 46.5, 44.2, 42.2, 36.9, 36.8, 35.6, 35.0, 34.4, 33.8, 30.9, 30.6, 28.7, 27.4, 26.5, 25.4, 23.5, 22.3, 17.3, 12.7. HR ESIMS m/z 389.2693 [M-H]^-^, C_24_H_37_O_4_ requires 389.2697.

#### 3-oxo-7β-hydroxy-5β-cholanoic acid (**3-oxoUDCA**)

An analytic sample was obtained by HPLC on a Nucleodur 100-5 C18 (5 μm; 4.6 mm i.d. x 250 mm), with MeOH/H_2_O (68:32) and trifluoroacetic acid 0.1% as eluent (flow rate 3 mL/min) (t_R_ = 8.44 min).

Selected ^1^H NMR (CD_3_OD, 400 MHz): δ_H_ 3.63 (1H, m), 2.53 (1H, t, *J* = 14.4 Hz), 2.42 (1H, m), 1.07 (3H, s), 0.97 (3H, d, *J* = 6.4 Hz), 0.73 (3H, s). ^13^C NMR (CDCl_3_, 100 MHz) δ 212.5, 179.3, 70.8, 55.6, 54.8, 44.3, 43.7, 43.2, 42.9, 39.9, 39.3, 36.9, 36.3, 36.1, 35.1, 34.3, 30.9, 30.7, 28.5, 26.7, 22.5, 21.6, 18.3, 12.1. HR ESIMS m/z 389.2695 [M-H]^-^, C_24_H_37_O_4_ requires 389.2697.

#### Synthesis of 3α-hydroxy-5α-Cholanoic Acid (**AlloLCA**), 3-oxo-5α-Cholanoic Acid (**3-oxoAlloLCA**), 3α-hydroxy-5-Cholenoic Acid (Δ^5,6^**-AlloLCA**), and 3β-hydroxy-5-Cholanoic Acid (**IsoAlloLCA**)

HDCA methyl ester (2.0 g, 4.91 mmol) was dissolved in N,N-dimethylformamide: pyridine anhydrous (2:1) and then imidazole and tert-butyldimethylsilyl chloride were added at 0°C. The mixture was stirred for 6 h at 0°C, then evaporated in vacuum and extracted with water and EtoAC (3 x 50 mL). The combined organic layers were dried on Na_2_SO_4_, filtered, and evaporated to furnish a crude residue that was purified on flash chromatography with 97:3 dichloromethane/methanol as eluent, furnishing 1.55 g of pure protected intermediate (2.97 mmol, 60% yield).

Pure protected product (1.55 g, 2.97 mmol) was dissolved in pyridine dry, and then p-toluenesulfonyl chloride (5 eq.) was added. The reaction was stirred at room temperature overnight. When the reaction was completed, the pyridine was removed under vacuum and the mixture extracted with H_2_O: EtOAc (3 x 50 mL). The combined organic layers were washed with NaHCO_3_ saturated solution and then with brine, dried on Na_2_SO_4_, filtered, and evaporated to furnish a C-6 monotosylated derivative **9** (1.9 g, 3.42 mmol, quantitative yield).

Monotosylated derivative **9** (1.9 g, 3.42 mmol) was then subjected to elimination of tosyl group at C6 for treatment with LiBr and Li_2_CO_3_, in DMF dry. The reaction was stirred overnight at reflux, then poured into water and extracted with ethyl acetate for three times (3 x 50 mL). The combined organic phases were washed with brine, dried with Na_2_SO_4_, filtered, and evaporated to give 1.7 g of derivative **10** as crude (3.48 mmol, quantitative yield). This product was solubilized in dry THF and then an excess of tetra-n-butylammonium fluoride (TBAF) solution (1M in THF dry) was added. When the reaction was completed, it was extracted for three times with H_2_O and EtOAc (3 x 30 mL). Then the organic layer was dried, filtered, and concentrated under vacuum to give intermediate **11** as crude residue in quantitative yield (1.5 g, 3.86 mmol).

A fraction of compound **11** (750 mg, 1.93 mmol) was dissolved in THF: MeOH anhydrous 1:1 v/v and then a small aliquot of palladium hydroxide was added. The reaction was stirred overnight under hydrogen pressure, to furnish an intermediate with a trans conformation between rings A and B (800 mg, 2.04 mmol, quantitative yield).

This compound hydrolyzed with NaOH pellets (10 eq.) in a solution of MeOH: H_2_O 1:1 v/v (20 mL) at reflux. When the reactions were completed, the resulting solutions were concentrated under vacuum, diluted with water, acidified with HCl 6 N and extracted with ethyl acetate (3 x 50 mL). The collected organic phases were washed with brine, dried over Na2SO4 anhydrous, filtered, and evaporated under reduced pressure to furnish **AlloLCA** (620 mg, 1.64 mmol), as crude residues, in 80% yield.

#### 3α-hydroxy-5α-Cholanoic Acid (**AlloLCA**)

An analytic sample was obtained by HPLC on a Nucleodur 100-5 C18 (5 μm; 4.6 mm i.d. x 250 mm), with MeOH/H_2_O (90:10) and trifluoroacetic acid 0.1% as eluent (flow rate 3 mL/min) (t_R_ = 15.3 min).

Selected ^1^H NMR (CDCl_3_, 400 MHz): δ_H_ 4.05 (1H, br s), 2.41 (1H, m), 2.26 (1H, m), 0.94 (3H, d, *J* = 6.4 Hz), 0.78 (3H, s), 0.66 (3H, s).

^13^C NMR (CDCl_3_, 100 MHz): δ_C_ 179.8, 66.7, 56.7, 55.9, 54.3, 42.6, 40.4, 39.7, 37.6, 35.8, 35.4, 35.3, 34.5, 32.2, 31.0, 30.9, 30.8, 28.2, 28.1, 24.1, 20.8, 18.2, 18.2, 12.1. HR ESIMS m/z 375.2901 [M-H]^-^, C_24_H_39_O_4_ requires 375.2905.

To a solution of **AlloLCA** (310 mg, 0.82 mmol) in acetone, 0.54 mL/mmol of Jones reagent solution (CrO_3_, H_2_SO_4_) as previously described.

#### 3-oxo-5α-Cholanoic Acid (**3-oxoAlloLCA**)

An analytic sample was obtained by HPLC on a Nucleodur 100-5 C18 (5 μm; 4.6 mm i.d. x 250 mm), with MeOH/H_2_O (90:10) and trifluoroacetic acid 0.1% as eluent (flow rate 3 mL/min) (t_R_ = 21.0 min).

Selected ^1^H NMR (CDCl_3_, 400 MHz): δ_H_ 2.45-1.05 (m, 29H), 0.95 (3H, d, *J* = 6.4 Hz), 0.74 (3H, s), 0.67 (3H, s).

^13^C NMR (CDCl_3_, 100 MHz): δ_C_ 207.1, 178.1, 56.7, 55.7, 51.5, 50.2, 42.4, 39.8, 39.7, 37.3, 35.3, 33.2, 31.9, 31.8, 31.0, 30.9, 30.7, 28.9, 28.1, 24.2, 20.8, 18.6, 18.3, 11.8. HR ESIMS m/z 373.2745 [M-H]^-^, C_24_H_37_O_3_ requires 373.2748.

A small aliquot of compound **11** (750 mg, 1.93 mmol) was subjected to hydrolysis in the same experimental condition previously described, to give **D**^5,6^**-AlloLCA** (440 mg, 1.17 mmol).

#### 3α-hydroxy-5-Cholenoic Acid (Δ^5,6^**-AlloLCA**)

An analytic sample was obtained by HPLC on a Nucleodur 100-5 C18 (5 μm; 4.6 mm i.d. x 250 mm), with MeOH/H_2_O (90:10) and trifluoroacetic acid 0.1% as eluent (flow rate 3 mL/min) (t_R_ = 16.2 min).

Selected ^1^H NMR (CDCl_3_, 400 MHz): δ_H_ 5.41(1H, m), 4.03 (1H, m), 2.58 (1H, m), 2.26 (1H, m), 2.41 (1H, m), 1.02 (3H, s), 0.95 (3H, d, *J* = 6.4 Hz), 0.69 (3H, s).

^13^C NMR (CDCl_3_, 100 MHz): δ_C_ 178.6, 138.4, 123.9, 67.2, 56.7, 55.7, 50.2, 42.3, 39.7, 39.6, 37.2, 35.3, 33.1, 31.9, 31.8, 30.8, 30.7, 28.8, 28.1, 24.2, 20.7, 18.6, 18.2, 11.8. HR ESIMS m/z 373.2743 [M-H]^-^, C_24_H_37_O_3_ requires 373.2748.

Starting from another aliquot of **HDCA methyl ester** (1 g, 2.46 mmol) as previously described ^2,4^ we obtained compound **12**, that was subjected to the hydrogenation and hydrolysis in the same experimental condition, above described.

#### 3β-hydroxy-5-Cholanoic Acid (**IsoAlloLCA**)

An analytic sample was obtained by HPLC on a Nucleodur 100-5 C18 (5 μm; 4.6 mm i.d. x 250 mm), with MeOH/H_2_O (90:10) and trifluoroacetic acid 0.1% as eluent (flow rate 3 mL/min) (t_R_=15.5 min).

Selected ^1^H NMR (CDCl_3_, 400 MHz): δ_H_ 3.60 (1H, m), 2.41 (1H, m), 2.28 (1H, m), 0.93 (3H, d, *J* = 6.5 Hz), 0.81 (3H, s), 0.66 (3H, s).

^13^C NMR (CDCl_3_, 100 MHz): δ_C_ 178.6, 71.4, 56.4, 55.8, 54.3, 44.8, 42.7, 40.0, 38.2, 37.0, 35.5, 35.4, 35.3, 32.0, 31.5, 30.9(2C), 30.8, 30.3, 28.7, 28.1, 21.2, 18.2, 12.1. HR ESIMS m/z 375.2903 [M-H]^-^, C_24_H_39_O_3_ requires 375.2905.

### Alpha Screen assay

Recombinant human LIFR (His Tag) and biotinylated recombinant human LIF were reconstituted as required by the manufacturer. Inhibition of LIFR/LIF binding by 35 BAs was measured by Alpha Screen (Amplified Luminescent Proximity Homogeneous Assay). The assay was carried out in white, low-volume, 384-well AlphaPlates (PerkinElmer, Waltham, MA, USA) using a final volume of 25 µL and an assay buffer containing 25 mM Hepes (pH 7.4), 100 mM NaCl, 0.05% (w:v) BSA and 0.005% Kathon. The concentration of DMSO in each well was maintained at 5%. LIFR (His Tag, final concentration 4.5 nM) was incubated with each bile acid (at 7 concentrations from 300 nM to 200 µM) or a vehicle for 45 min under continuous shaking. Then, LIF was added (biotinylated, final concentration 9 nM), and the samples were incubated for 15 min prior to adding His-Tag acceptor beads (final concentration 20 ng/µL) for 30 min. Then, streptavidin donor beads were added (final concentration 20 ng/µL), and the plate was incubated in the dark for 2 h and then read in an EnSpire Alpha multimode plate reader (PerkinElmer, Waltham, MA, USA).

### Cell cultures

#### 2D CELL LINES

Experiments were conducted using various gastrointestinal cell lines, with a focus on the human gastric cancer cell line, MKN45, which was cultured in RPMI 1640 medium supplemented with 10% Fetal Bovine Serum (FBS), 1% L-Glutamine, and 1% Penicillin/Streptomycin. HepG2, an immortalized human hepatocarcinoma cell line, was cultured in E-MEM with 10% FBS, 1% L-Glutamine, and 1% Penicillin/Streptomycin. The human pancreatic cell line, MIA-PaCa-2, and the human intestinal epithelial cell line, Caco2, were cultured in D-MEM containing 10% FBS, 1% L-glutamine, and 1% penicillin/streptomycin.

#### 3D CELL LINES. Gastric glands extraction

Human gastric glands were extracted from neoplastic mucosa excided from gastric cancer patients. Gastric mucosa resection was obtained from 8 patients undergoing surgical resection at the Surgery Unit of the Perugia University Hospital (Italy). Informed written consent was obtained from each patient before surgery. None of the patients had received chemotherapy or radiation before surgery. (permit FI00001, n. 2266/2014 and permit FIO0003 n.36348/020).

Murine gastric glands were isolated from the antrum of 4-8 weeks C57BL6/J mice. Mice were housed under regulated temperature (22°C) and photoperiods (12:12-h light/dark cycle), allowed unrestricted access to standard mouse chow and tap water. The general health of the animals was monitored daily by the Veterinarian in the animal facility (permission n. 309-2022-PR). **Organoids establishment**. Human mucosa and murine stomach tissue was washed in cold PBS supplemented with Primocin. Then, tissue was cut in small fragments and incubated in Stripping buffer (HBSS with 10% FBS, 25 mM Hepes and 5 Mm EDTA) for 20 min at 37 °C with shaking. The fragments were subjected to enzymatic digestion by collagenase (1,5 mg/mL) and hyaluronidase (20 µg/mL) in 10 mL Advanced DMEM F12 for 1 h at 37 °C with shaking. The gland suspensions were passed through 100 µM filter and washed twice in Advanced DMEM F12, seeded into Matrigel (50.000/50μL) and overlaid with Advanced DMEM F12 medium containing HEPES, L-Glutamine, Primocin, B27, N2, n-Acetylcysteine 1 mM, EGF (50 ng/mL), R-spondin1 (200 ng/mL), Noggin (200 ng/mL), Wnt (200 ng/mL), FGF10 (200 ng/mL), Gastrin (10 nM), TGFβ-inhibitor (A 83-01 0.5 mM), RHOK-inhibitor (Y-27632 10 µM) and LIF (10 ng/mL).

### Transactivation Assay

STAT3 transactivation was performed on HepG2 as described previously ^5^. On day 0, cell were seeded at 7.5 × 104 cells/well, on day 1, cells were transiently transfected with the reporter plasmid pGL4.47[luc2P/SIE/Hygro] (200 ng), a vector encoding the hLIFR (100 ng) and CD130 (IL6ST) (100 ng), and finally a vector encoding the human RENILLA luciferase gene (pGL4.70) (100 ng). On day 2, cells were exposed to the LIF (10 ng/mL) alone or in combination with DCA, TDCA, GDCA, 3-oxoDCA, LCA, TLCA, GLCA, 3-oxoLCA (1, 3, 10 and 20 μM). Then, after 24 h, the cells were lysed in 100 μL of lysis buffer (25 mM Tris-phosphate, pH 7.8; 2 mM dithiothreitol (DTT); 10% glycerol; 1% Triton X-100). Then, 10 μL cellular lysates were assayed for luciferase and RENILLA activities using the Dual-Luciferase Reporter assay system. Luminescence was measured using a Glomax 20/20 luminometer. LUCIFERASE activities (RLU) were normalized with RENILLA activities (RRU).

### Protein and Ligand preparation

The three-dimensional (3D) X-Ray structure (PDB ID: 3E0G)^7^ of the human LIFR (hLIFR) (Uniprot ID Code: P42702) was retrieved from the RCSB Protein Data Bank and subjected to the Maestro’s Protein Preparation Wizard tool (Schrödinger Release 2022-4) in order to assign bond orders, add hydrogen atoms, adjust disulphide bonds, add caps to chains break, and assign residues protonation state at pH 7.4 with the proPKa module. The natural bile acids (BAs) library was prepared using the LigPrep (LigPrep. Schrödinger, release 2022–4, LigPrep; Schrödinger, LLC: New York, NY, USA, 2022) and Epik (Schrödinger; Release 2022-4: Epik, S., LLC, New York, NY, USA, 2022) modules to generate and optimize the 3D structures of the ligands at the protonation states of physiological pH 7.4 [1].

### Docking

The above prepared 3D structure of *h*LIFR was used to perform two steps docking protocol already successfully adopted in two our previous works ^5,9^: *i)* the first step was performed with the QM-Polarized Ligands Docking (QPLD) (Schrödinger Release 2021-4) algorithm in order to improve the docking accuracy by considering ligand charges derived from ab-initio calculations (Glide, Schrödinger, LLC, New York, NY, 2021; Jaguar, Schrödinger, LLC, New York, NY, 2021; QSite, Schrödinger, LLC, New York, NY, 2021.); *ii)* the most energetically favorable protein-ligand poses were, then, submitted to the second Induced-Fit Docking (IFD) protocol (Glide, S., LLC, New York, NY, USA, 2021; Prime, S., LLC, New York, NY, USA, 2021), in order to predict the concomitant effect of ligand docking on the protein structure (Table S1). Briefly, the centroid of the hLIFR ligand binding site, delimited by L2 and L3 loops was used to generate the inner grid box coordinate (10.0 Å size). Ten docking poses were saved for each ligand in the QPLD step, and the most energetically favorable poses were sent to IFD second step procedure, using the extended sampling protocol which generates A maximum of 80 poses in an energy window for the ligand conformational sampling equal to 2.5 kcal/mol.

### Molecular Dynamics simulations (MDs)

The best scored IFD docking pose of DCA, gDCA, tDCA, 3-oxo-DCA, 3-oxo-LCA and tLCA BAs were submitted to 150 ns of MDs using CUDA version of the AMBER22 package ^11^ Each complex was prepared using the LEaP module of AmberTools22. Specifically, protein was treated with the using the Amber ff14SB force field ^13^, while ligand charges were, instead, calculated using the restrained electrostatic potential (RESP) fitting procedure ^15^. Firstly, the Gaussian16 package ^17^ was used to calculate the ligand ESP using the 6-31G* at the Hartree-Fock (HF) level of theory. Then, RESP charges were retrieved using the Antechamber module implemented in AmberTools22 package ^19^, coupled with the general amber force field (GAFF2) parameters ^21^. Each system was immersed in a 10 Å layer cubic water box using the TIP3P water model parameters ^23^ and then neutralized by adding Na+ and Cl-ions. A cut-off of 8 Å was used for non-bonded short-range interactions, while long-range electrostatic interactions were computed by means of the Particle Mesh Ewald (PME) method using a 1.0 Å grid spacing in periodic boundary conditions. The SHAKE algorithm was applied to constraint bonds involving hydrogen atoms, with a 2 fs integration time step. Each system was firstly minimized in four steps as described in our previous works ^5,9^ and successively, water molecules thermally equilibrated as previously described ^5,9^. Trajectories and data were processed and analyzed using the CPPTRAJ module ^25^ and the Visual Molecular Dynamics (VMD) graphics ver. 1.9.3 ^27^. For the most representative cluster population, intermolecular interaction energy was analysed via the Molecular Mechanics/Generalized Born Surface Area (MM/GBSA) equation ^29^ (Supporting information Tables S2 and S3). All images were rendered using Maestro GUI Suite 2022-4 (Schrödinger Release 2022-4) and Adobe Illustrator (Adobe Systems, San Jose, CA, USA).

### Co-Immune precipitation (Co-IP)

Co-IP was performed on MKN45 proteins using Abcam’s Immunoprecipitation Kit. 1.5 *10^6^ of MKN45 cells were exposed to LIF (10 ng/mL) alone or in combination with GDCA (3 µM) for 1 hour. Cells were then washed 1 time with ice-cold PBS, scraped and lysed according to the manufacturing instruction. 300 µg total proteins were pre-cleared on a rotating wheel for 1 h at 4°C using protein A/G Sepharose beads (kit provided). Immunoprecipitation was performed overnight at 4°C with the followings antibodies: 3 µg anti-LIFR antibody or 1 µg anti-IgG used as a negative control in the presence of 25 µL of protein A Sepharose (kit provided). The resultant immunoprecipitates were washed three times with 1 mL of wash buffer and resuspended in 25 µL of Tris-Glycine SDS Sample buffer 2X. Anti-LIFR immunoprecipitates were used for western blotting using the antibodies anti-LIFR and anti-gp130.

### Cell proliferation Assay

The cell viability assay was done using the CellTiter 96 Aqueous One Solution Cell Proliferation Assay, a colorimetric method for accessing the number of viable cells in proliferation proliferation as described previously ^5^. MKN45 cells were seeded at 36 *10^3^ cells/100 uL well into 96-well tissue culture plate. After 24 h, cells were serum starved for 24 h. Cells were exposed to LIF alone or in combination with 1, 3, 10 and 20 μM of DCA, TDCA, GDCA, 3-oxoDCA, LCA, TLCA, GLCA, 3-oxoLCA. Similarly, MIA PaCa-2, HepG2 and Caco2 were treated with LIF or DCA (3 μM), TDCA (10 μM), GDCA (3 μM), 3-oxoDCA (3 μM), LCA (10 μM), TLCA (10 μM), GLCA (10 μM), 3-oxoLCA (10 μM). Then cell proliferation was assessed as mentioned above. Absorbance was measured using a 96 well reader spectrophotometer (490 nm). In these experiments each experimental setting was replicated ten folds. For analysis the background readings with the medium alone, were subtracted from the samples read-outs.

### Flow-cytometry

MKN45 cells were seeded in 6-well tissue culture plate (cell density 700 × 10^3^/well) and cultured as specified above. Cells were serum-starved for 8 h and then incubated with LIF (10 ng/mL) alone or plus DCA (3 μM), TDCA (10 μM), GDCA (3 μM), 3-oxoDCA (3 μM), LCA (10 μM), TLCA (10 μM), GLCA (10 μM), 3-oxoLCA (10 μM) or a vehicle for 24 h. The intracellular flow cytometry staining for Ki-67 was performed using the following reagents: Ki-67 Monoclonal Antibody and 7-AAD to characterize the cell cycle phases G0-G1 and S-G2-M. Before intracellular IC-FACS, staining cells were fixed for 30 min in the dark using IC Fixation buffer and then permeabilized using Permeabilization buffer (10X). The staining for Annexin V was performed using the Annexin V Antibody to evaluate the apoptosis rate. Briefly, 5 μL of Annexin V. Antibody was added to each 100 μL of cell suspension, and cells were incubated the at room temperature for 15 min.

Cells were analyzed with FACS Fortessa. Data was analyzed with FlowJo software and the gates set using a fluorescence minus-one (FMO) control strategy. FMO controls are samples that include all conjugated Abs present in the test samples except for one. The channel in which the conjugated Ab is missing is the one for which the fluorescence minus one provides a gating control.

### RNA extraction

RNA was extracted from MKN45 cell lines and human patient derived oranoids (hPDOs), was extracted using The kit Direct-zol™ RNA MicroPrep w/ Zymo-Spin™ IIC Columns. RNA extracted was used for qPCR analysis.

### Reverse transcription of mRNA and Real time (RT)-PCR

After purification from genomic DNA using DNase I, 1 μg of RNA was reverse transcribed using the OptiFast cDNA Synthesis Kit in a 20-μL reaction volume; 10 ng of cDNA was amplified in a 20-μL solution containing 200 nM each primer and 10 μL of SYBR Select Master Mix. All reactions were performed in triplicate using the following thermal cycling conditions: 3 min at 95 °C, followed by 40 cycles of 95 °C for 15 s, 56 °C for 20 s, and 72 °C for 30 s, using a StepOnePlus system. The relative mRNA expression was calculated accordingly to the ΔCt method. Primers were designed using the software PRIMER3 (http://frodo.wi.mit.edu/primer3/) using published data obtained from the NCBI database. The primers used for mouse genes were as following (forward and reverse):

hCMYC (for TTTCGGGTAGTGGAAAACCA; rev CACCGAGTCGTAGTCGAGGT).

hSNAIL1 (for ACCCACACTGGCGAGAAG; rev TGACATCTGAGTGGGTCTGG);

hVIM (for TCAGAGAGAGGAAGCCGAAA; rev ATTCCACTTTGCGTTCAAGG);

hBCL2 (for GAAACTTGACAGAGGATCATGC; rev TCTTTATTTCATGAGGCACGTT);

hLIFR (for GCTCGTAAAATTAGTGACCCACA; rev GCACATTCCAAGGGCATATC);

hLIF (for CCCTGTCGCTCTCTAAGCAC; rev GGGATGGACAGATGGACAAC);

### Western Blot Analysis

MKN45 were lysed in RIPA lysis buffer containing phosphatase and protease inhibitors cocktail; aliquots from each sample containing 50 µg of protein were separated on Novex WedgeWell 10% Tris-Glycine gel (Invitrogen) and transferred to nitrocellulose membrane with iBlot 2 Dry Blotting System (Invitrogen).The blots were subsequently blocked for 1 h with 5% milk powder in Tris-buffered saline (TBS)/Tween 20 at RT and then probed overnight (at 4 °C) with primary antibodies against GAPDH (1:1000), STAT3 (1:1000), pSTAT3 (1:1000). After overnight incubation, anti-rabbit IgG and anti-mouse horseradish peroxidase-labeled secondary antibody, at a dilution of 1:1000, were used. Positive signals were developed by Immobilon Western Chemiluminescent HRP Substrate (Merck Millipore) and the images was achieved with iBright Imaging Systems (ThermoFisher). Quantitative densitometry analysis was performed using ImageJ Software. The degree of STAT3 phosphorylation was calculated as the ratio between the densitometry readings of GAPDH and p-STAT3/ STAT3.

### UPLC-MSMS analysis of 35 BAs

GC biopsies were preserved at −80°C and lyophilized. Then, 20 mg of each sample was manually homogenized using a potter pestle and dissolved in MeOH at a final concentration 100 µg/µL for an opportune extraction for 2 h. Finally, they were diluted 5 times in a solution made of 50% H2O/50% MeOH, 0.1% formic acid (FA) and 5 mM ammonium acetate (AmAc).

Stock solutions of the individual BAs were distinctly prepared in MeOH, mixed and diluted in 50% H2O/50% MeOH, 5 mM AmAc and 0.1% FA to obtain the calibration standards ranging from 25 nM to 400 nM.

UHPLC-MRM-MS analyses were performed on a QTRAP 6500 (AB Sciex) equipped Shimadzu Nexera LC and Auto Sampler systems. A mixture of 35 BAs was detached on a Luna Omega 1.6 μm Polar (C18, 100 Å, 50 × 2.1 mm; Phenomenex) at 40°C, and at a flow rate of 400 μl/min. Mobile phase A was H2O, 5 mM AmAc, 0.1% FA, and mobile phase B was MeOH, 5 mM AmAc, 0.1% FA. The gradient started at 50% B, increased to 55% B in 3.5 min and then to 95% B in 19.5 min, was kept at 95% B for 1 min and then decreased to 50% B and was kept to re-equilibrate for 6 min. Q-TRAP 6500 was operated in negative MRM scanning mode with the following parameters:

Negative mode: declustering potential (DP) at −150 V, entrance potential (EP) at −12 V, collision energy (CE) at −15V and cell exit potential (CXP) at −30 V. Curtain gas was set at 30, ion source gas 1 and 2 at 25 and ion spray voltage at −4500. Resolution Q1 and Q2 was unitary.

For quantification, the extracts were injected alongside the calibration mixtures. The area of each peak was measured through the Analyst software (ABSciex) using the following mock transitions:

t HCA at 5.61 min, mock MRM at m/z of 514; tCA at 7.54 min, mock MRM at m/z of 514; tUDCA at 5.54 min, mock MRM at m/z of 498; tHDCA at 6.51 min, mock MRM at m/z of 498; tCDCA at 10.18 min, mock MRM at m/z of 498; tDCA at 10.75 min, mock MRM at m/z of 498; tLCA at 12.82 min, mock MRM at m/z of 482; HCA at 11.53 min, mock MRM at m/z of 407; CA at 12.71 min, mock MRM at m/z of 407; UDCA at 11.31 min, mock MRM at m/z of 391; HDCA at 1.37 min, mock MRM at m/z of 391; CDCA at 15.25 min, mock MRM at m/z of 391; DCA at 15.55 min, mock MRM at m/z of 391; LCA at 17.66 min, mock MRM at m/z of 375; isoalloLCA at 16.83 min, mock MRM at m/z of 375; alloLCA at 18.04 min, mock MRM at m/z of 375; Δ5,6LCA at 17.21 min, mock MRM at m/z of 373; 3-oxo-LCA at 17.33 min, mock MRM at m/z of 373; 3-oxo-allo-LCA at 17.99 min, mock MRM at m/z of 373; 3-oxo-UDCA at 10.97 min, mock MRM at m/z of 389; 7k-LCA at 12.02 min, mock MRM at m/z of 389; 3-oxo-CDCA at 13.94 min, mock MRM at m/z of 389; 3-oxo-DCA at 14.11 min, mock MRM at m/z of 389; 3-oxo-CA at 10.81 min, mock MRM at m/z of 405; t7kLCA at 6.05 min, mock MRM at m/z of 496; t3-oxo-LCA at 11.85 min, mock MRM at m/z of 480; gHCA at 8.05 min, mock MRM at m/z of 464; gCA at 9.87 min, mock MRM at m/z of 464; g-3-oxo-LCA at 14.23 min, mock MRM at m/z of 430; gLCA at 14.97 min, mock MRM at m/z of 432; g7kLCA at 8.78 min, mock MRM at m/z of 446; gUDCA at 7.94 min, mock MRM at m/z of 448; gHDCA at 8.93 min, mock MRM at m/z of 448; gCDCA at 12.35 min, mock MRM at m/z of 448; gDCA at 13.01 min, mock MRM at m/z of 448.

### Histological techniques

#### Immunofluorescence analysis (IF)

Immunofluorescence staining was performed on murine gastric organoids and hPDOs derived from gastric cancer resections.

After removing the culture medium, organoids were washed rapidly once with PBS (1X) and then fixed in 4% PFA for 20 minutes at room temperature (RT). After fixation, 4% PFA was removed and organoids were washed gently with rocking, using PBS for three times, each wash lasting for 5 minutes. Subsequently, the organoids were permeabilized with PBS containing 0,5% Triton for 15 minutes at RT, followed by washing, with gently rocking in an IF Buffer (PBS+ 0,2% Triton X-100 + 0,05% Tween-20) three times, each for 5 minutes. Finally, organoids were incubated for 1 hat RT using an IF buffer containing 2,5% BSA.

Primary antibodies anti-LIFR (1:100) and anti-E-cadh (1:100) were diluted in IF buffer + 1% BSA and incubated overnight at 4° C. The next day, primary antibodies were recovered and organoids were washed, with rocking, in IF buffer three times. Subsequently, organoids were incubated at RT for 2 h with secondary antibodies: Goat Anti-Rabbit IgG H&L Alexa Fluor® 488 and Goat Anti-Rat IgG (H+L) Alexa Fluor® plus 568 diluted in IF buffer with 1% BSA. After another set of three 5 minute washes with IF buffer, nuclei were labelled with DAPI and incubated for 5 minutes at RT. Following the last 5-minute wash with IF buffer, the organoids were ready for acquisition using the Nikon Eclipse Ti Confocal Spinning Disc CrestV2 or could be stored at 4°C.

#### Hematoxylin and Eosin (H&E)

For histological examination, murine gastric organoids were fixed in 10% formalin, embedded in paraffin and then sectioned. Sections were then stained with Hematoxylin/Eosin (H&E), for morphometric analysis.

### Quantification and statistical analysis

For comparisons involving more than two groups, we employed either a one-way ANOVA or a paired Student t-test with Welch’s correction for comparisons of two groups, as appropriate (*p < 0.05) using GraphPad Prism 8.0 software.

Before conducting correlation studies, we assessed the normality of the data using the Kolmogorov-Smirnov test (p < 0.05). Correlation coefficients were calculated using Pearson’s r for datasets with a Gaussian distribution and Spearman’s rank correlation coefficient (Spearman’s rho) for datasets that did not follow a Gaussian distribution.

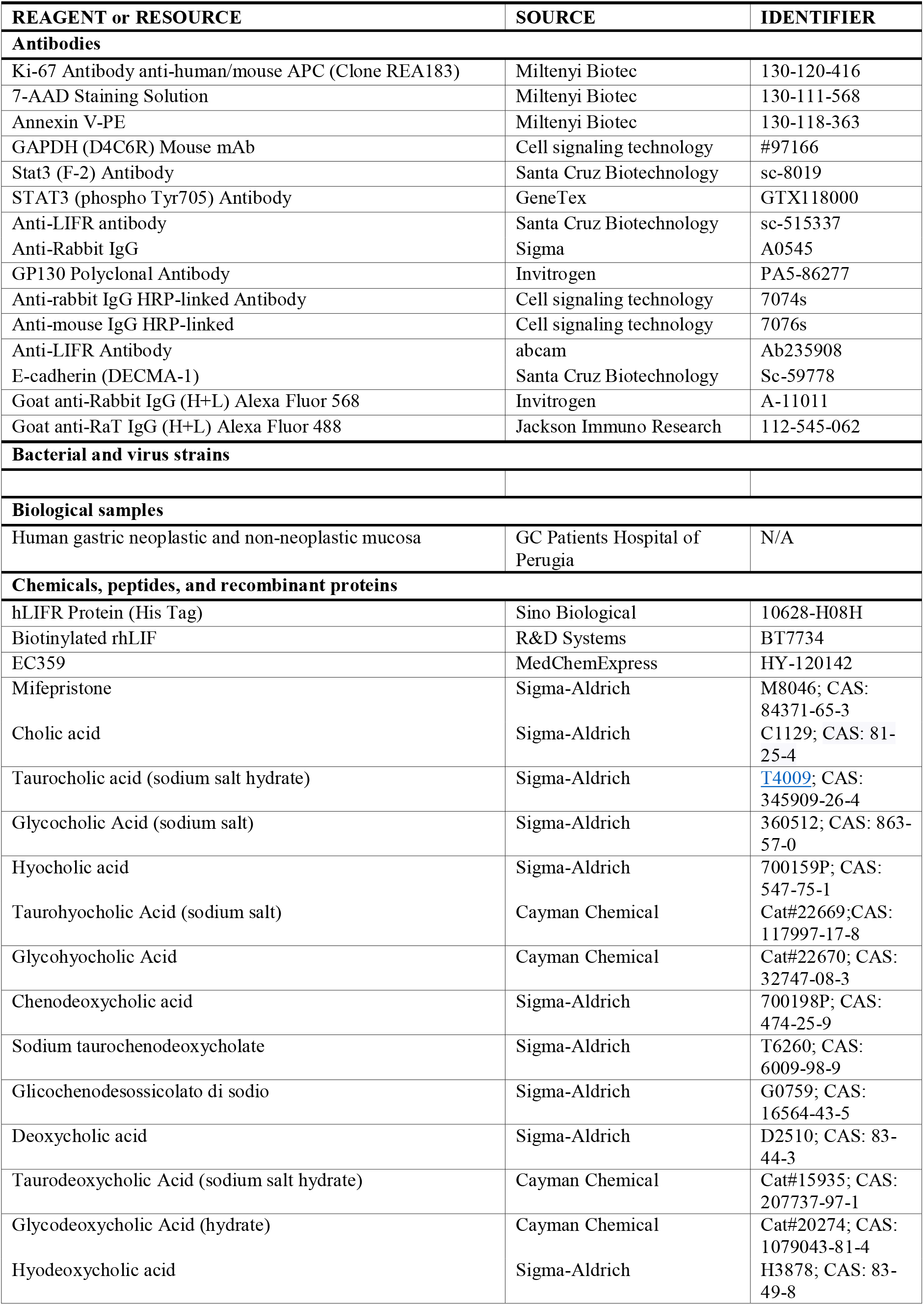
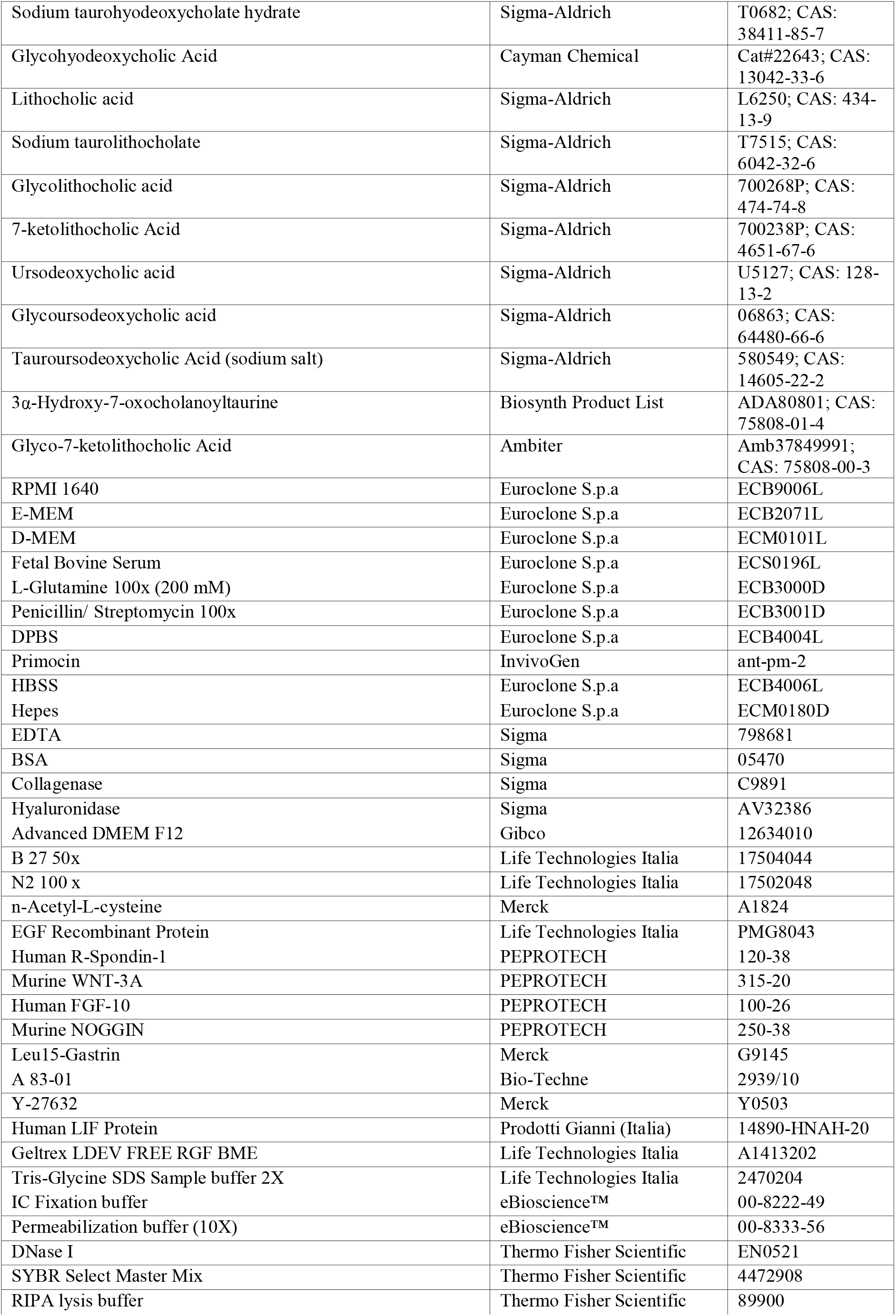
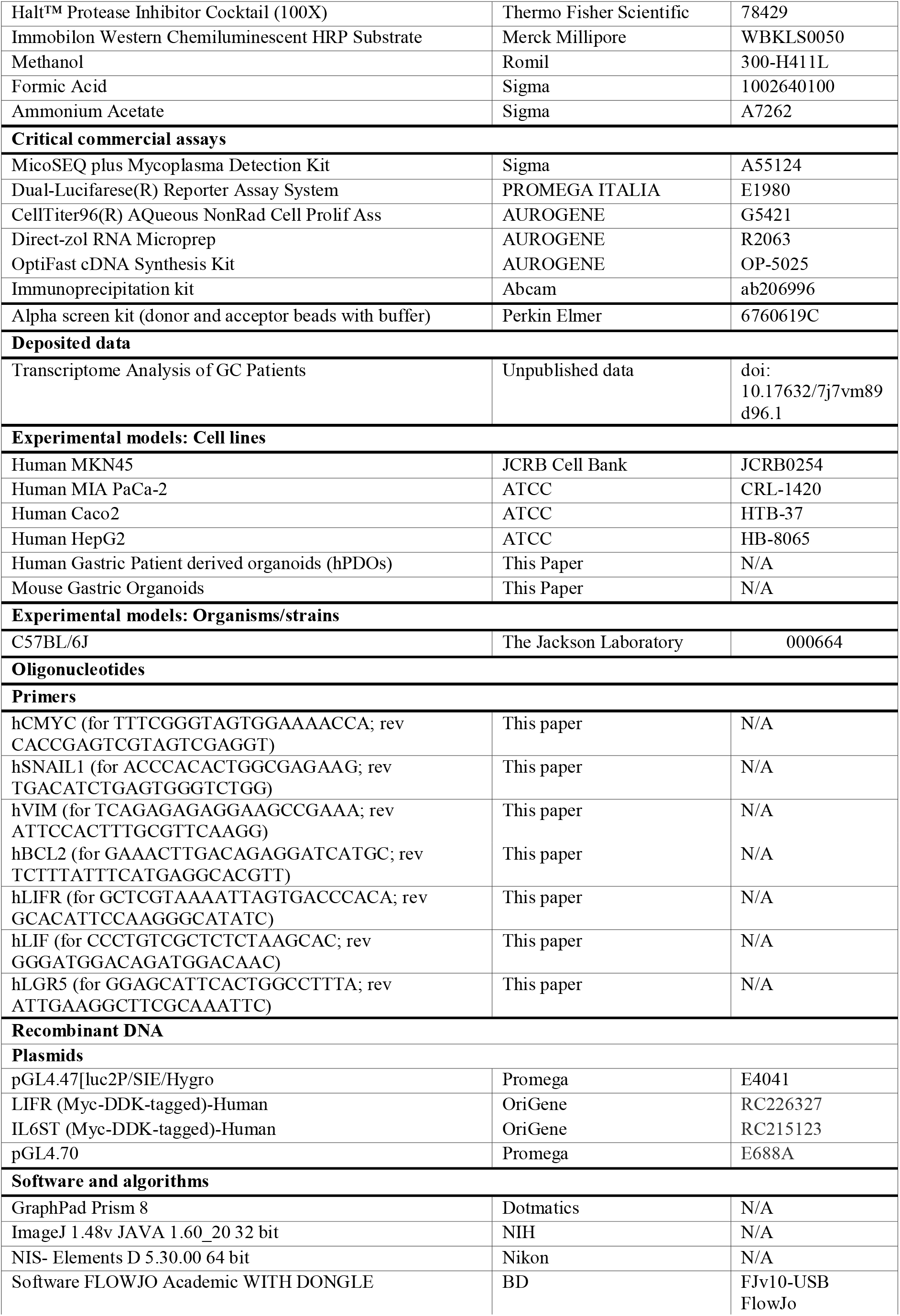
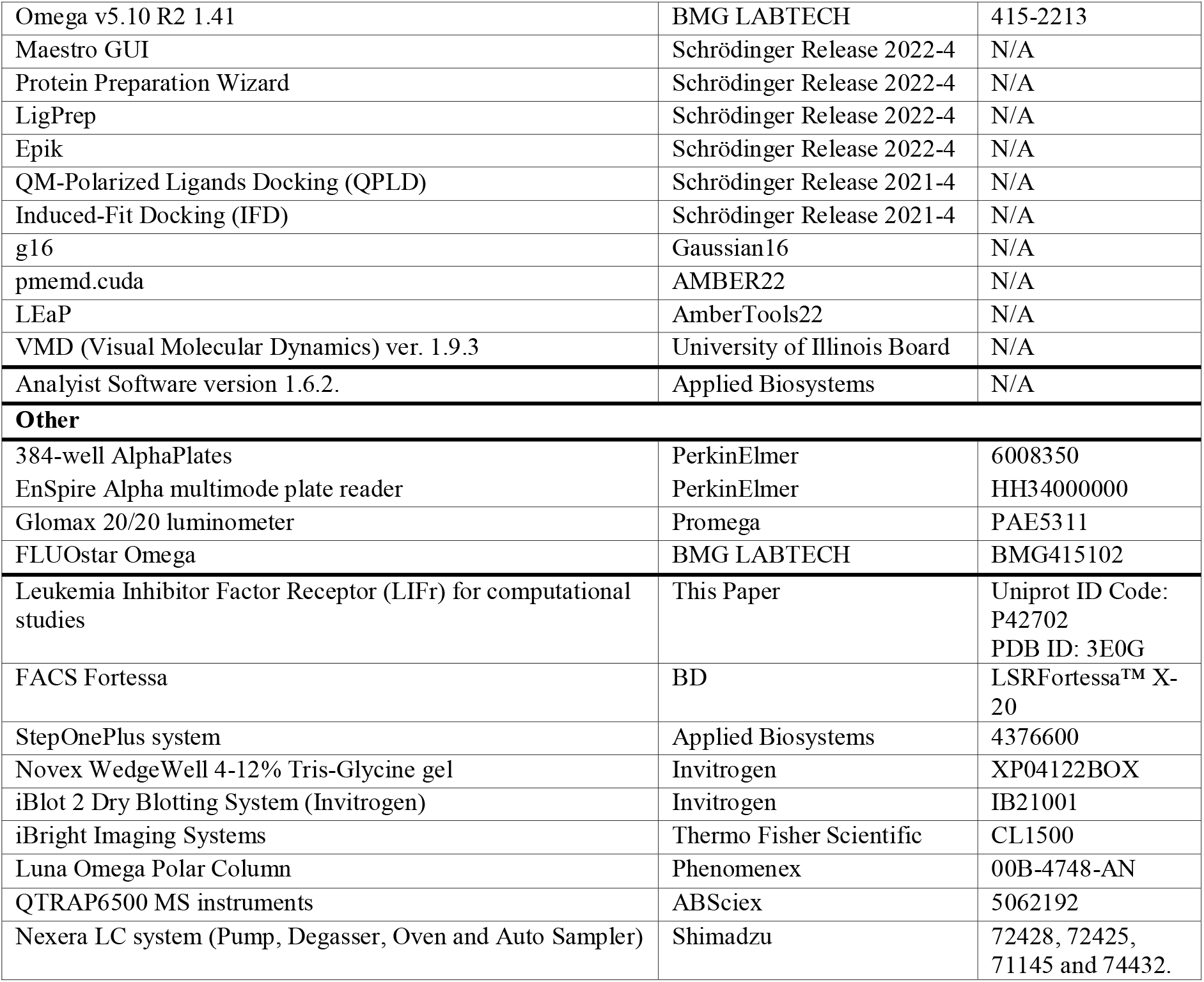

## Results

In this study, 35 different bile acids, including primary and secondary bile acids, their oxo-derivatives and some conjugated forms were tested. While 24 compounds were purchased from Sigma-Aldrich, 11 different derivatives (G-3-oxoLCA, T-3-oxoLCA, 3-oxoUDCA, 3-oxo-AlloLCA, and Δ5,6-LCA, 3-oxoLCA, 3-oxoCDCA, 3-oxoCA, 3-oxoDCA, AlloLCA, and IsoAlloLCA, Figure S1) of this large library of compounds were synthetized and reported here in Figure 1 A-B. Full details of the synthesis are given in the experimental section.

**Figure 1.**
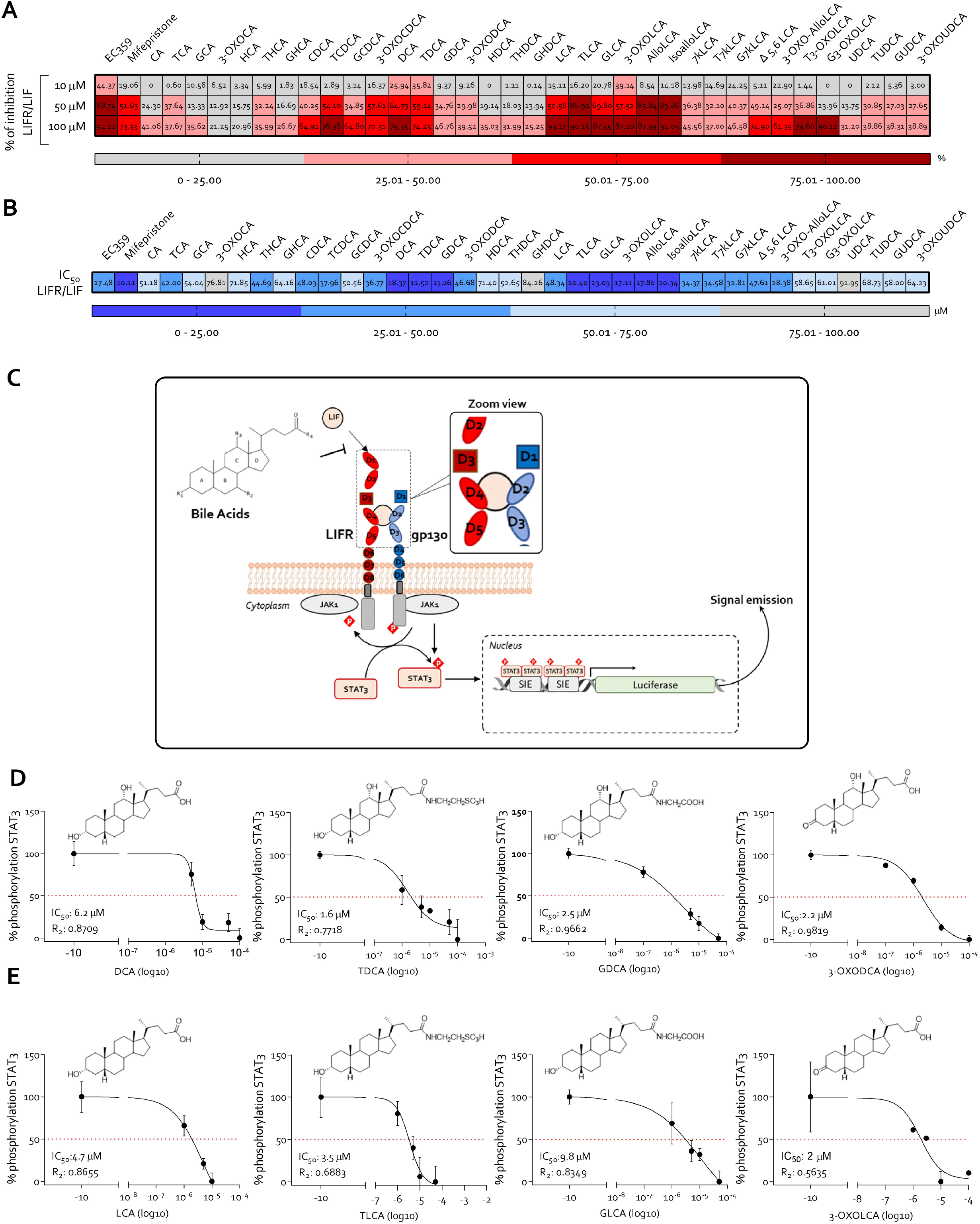
Bile acids are antagonists of LIF/LIFR interaction. The potential inhibitory of bile acids as antagonists of LIF/LIFR was investigated using a cell-free AlphaScreen assay. hLIFR was incubated with bile acids and then LIF capacity binding was measured. Panel **A)** shows bile acids percentage of LIF/LIFR inhibition at the concentration of 10-50 and 100 μM (from top to bottom). Red squares represent higher % of inhibition while grey ones represent the lower. Panel **B)** provides a detailed analysis of IC50 values of bile acids. Blue squares represent the higher % of inhibition, while grey squares represent the lower. Panel **C)** illustrates the schematic strategy of transactivation assay of STAT3 detailed in Material and method section. STAT3 transactivation on HepG2 cells of **D)** DCA, TDCA, GDCA and 3-oxoDCA; **E)** LCA and its T; G and 3-oxo derivatives. (* represents statistical significance versus NT, and # versus LIF, *p < 0.05).

To examine whether primary and secondary bile acids modulate the binding between LIF and LIFR, we have first settled up a cell free assay based on an Alpha Screen platform ^9^. Because natural bile acids activate the majority of their receptors in the 1-100 µM range^55^, the assays were carried out using this concentrations range and the potency of various bile acids in inhibiting LIF/LIFR interaction (IC50) was compared to that of two well characterized LIFR antagonists, EC359^56^ and mifepristone^5^. The results of these studies, shown in Figure 1A, demonstrated that, while among 35 different bile acids, only DCA, TDCA and 3-oxoLCA effectively inhibited LIF/LIFR interaction at 10 µM, the number of potential LIF/LIFR antagonists rose significantly at 50 µM, and almost all bile acids, except 3-oxoCA and HCA, prevented LIF/LIFR interaction at the concentration of 100 µM.At the concentration of 50 µM, CDCA, DCA, LCA and their Glyco (G), Tauro (T) and 3-oxo derivatives reduced LIFR activation by > 50%, and the inhibition rose to ≈80% at 100 µM.

The calculated IC50 for DCA and LCA and some of their derivatives was in the 10-20 µM range (Figure 1B). Thus, LIF/LIFR inhibition by various bile acids occurs in the same range of concentration as required for activation of membrane and nuclear receptors such as S1PR2, FXR and GPBAR1^55^. These results demonstrated that bile acids might function as LIF/LIFR binding antagonists at physiological tissue concentrations (Figure 1B).

Previous studies have shown that binding of LIF to LIFR induces the assembly of a LIFR/gp130 heterodimer that recruits and activates the Janus kinase 2 (JAK2), which in turn phosphorylates STAT3^43^. Once phosphorylated, STAT3 dimerizes and translocates to the nucleus binding to STAT3 inducing elements (SIE) and the promoter of target genes. To test whether bile acids interfere with the signalling pathway, we challenged HepG2 cells, a hepatic cancer cell line, transfected with viral vectors encoding for hLIFR and gp130 and STAT3 responsive elements cloned upstream the *Renilla luciferase* gene^5^, with DCA and LCA. The results of these experiments, shown in Figure 1 D-E, confirmed that DCA and LCA and their T, G and the corresponding 3-oxo derivatives effectively inhibited STAT3 phosphorylation induced by LIF at concentrations < 10 µM. DCA and its derivatives were more potent than LCA analogues in abrogating LIF-induced phosphorylation and their respective IC_50_s were as follow: DCA ≈ 6.2 µM, TDCA ≈ 1.6 µM, GDCA ≈ 2.5 µM and 3-oxoDCA ≈ 2.2 µM (Figure 1 D). Similarly, LCA and its derivatives reduced STAT3 phosphorylation though at slightly higher concentrations: LCA ≈ 4.7 µM, TLCA ≈ 3.5 µM, GLCA ≈ 9.8 µM and 3-oxoLCA ≈ 2 µM (Figure 1 E). Thus, secondary bile acids inhibit LIF-induced STAT3 phosphorylation at the same concentrations (1-2 µM) required for GPBAR1 activation^20^.

To disclose the molecular basis of LIF/LIFR inhibition by secondary bile acids, we have then simulated the binding mode of DCA and its derivatives GDCA, TDCA and 3-oxoDCA and of two most active LCA derivatives, TLCA and 3-oxoLCA. We applied the same protocol used in our previous studies on LIFR inhibitors ^5,9^ consisting of two steps of docking calculations, better discussed in the Methods section, followed by 150 ns of Molecular Dynamics simulations (MDs). Docking calculations were addressed in the previously predicted pocket on the Ig-like domain D3, defined by the loops L2 and L3 involved in the LIF binding (Figure 2A). During 150 ns of MDs of the best docking poses, we observed a high flexibility mainly of the L3 loop, as already revealed for other inhibitors ^5,9^, which induces a continuous change in the shape of the binding site that affected also the binding stability of some ligands. Indeed, while compounds DCA, 3-oxoDCA, 3-oxoLCA and TDCA showed a stable binding inside the two loops L2 and L3, GDCA and TLCA demonstrated a less stable binding mode, as retrieved by the Ligand Root Mean Square Deviation (L-RMSD) plot in Supporting information Figure S2A. Nonetheless, the most populated clusters of the most potent inhibitors analysed, TLCA, TDCA and 3-oxoLCA (Supporting Information Figure S2B and Table S2), shared similar interacting features (Supporting Information Figure S3). Specifically, a salt-bridge between the sulfonic acid group and the K332 (loop L3) was retrieved in both the tauro conjugates (TDCA and TLCA) and between the glutamic acid group of 3-oxoLCA and R330 (loop L3). In addition, the rings C and D of the steroidal scaffold of TLCA, 3-oxoLCA and TDCA are sandwiched between Y342 of loop L3 and Y318 located in the β-sheet of the D3 domain, with the oxygen atom in position 3 engaging H-bonds interactions with the backbone of L2 loop residues R306, T308 and L310, respectively (Supporting information, Figures S3). The milder inhibitor DCA, during MDs, established a stable H-bond with T308, but its carboxylic group in the side chain did not show any stable interaction (Supporting information, Figures S3).

**Figure 2.**
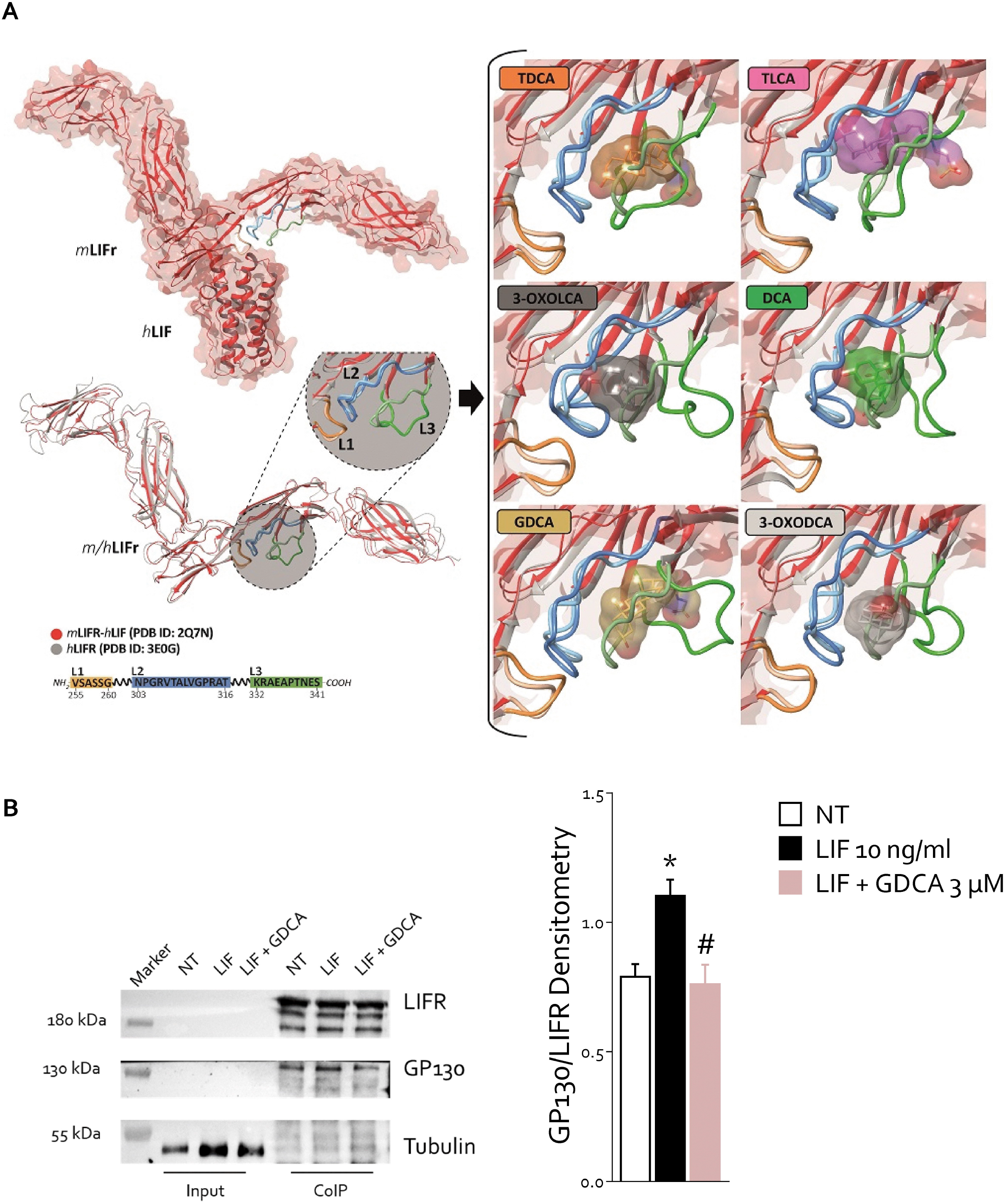
Bile acids inhibit LIFRβ:gp130 heterodimer. **A)** An overview of both hLIFR-h/mLIF complex and all BAs into the binding site. The protein backbone is displayed in red ribbon, while in the zoom view, the binding site is defined by loops L1 (255-VSASSG-260), L2 (303-NPGRVTALVGPRAT-316), and L3 (332-KRAEAPTNES-341) are highlighted in orange, blue, and green, respectively. On the right, the BAs poses are visualized and superimposed to the mLIFR– hLIF complex. **B)** Representative co-Immune precipitation analysis of LIFR and gp130 (on left) and densitometric analysis demonstrating gp130/LIFR ratio (on right). (* represents statistical significance versus NT, and # versus LIF, *p < 0.05).

Interestingly, both less potent LIFR inhibitors analysed, GDCA and 3-oxoDCA, did not show any stable interaction between the functional groups in position 3 and residues in the loop L2, but only with residues bearing to loop L3. Additional polar and hydrophobic contacts contributed to further stabilize the interaction of bile acids with L2 or L3. In particular in the DCA series, the hydroxyl group at position 7 engaged an additional H-bond with T316 (in DCA, TDCA and GDCA) or with T338 (in 3-oxoDCA). Overall, the computational studies suggested that all ligands analysed behaved as a wedge, separating the L2 and L3 loops, altering the conformation of the loops L2 and L3, which are highly involved in binding with LIF (Figure 2A).

Because the assembly of LIFR:gp130 complex is essential for LIFR signalling ^7^ (Figure 1C), we then investigated whether bile acids impact on the formation of LIF/LIFR complex. For these studies, we have used GDCA since it is the most represented of LIFR antagonists detected in tumour (see below) and in MKN45 cells, a GC cell line. The results of the co-immunoprecipitation (Co-IP) studies (Figure 2 B) demonstrated that, while exposure of MKN45 cells to LIF 10 ng/mL, promotes the assembly of the LIFR/gp130 complex (Figure 2B, lane 6), this pattern has been, reversed by co-treating the cells with 3 µM of GDCA (Figure 2B lane 7). Together these results demonstrated that secondary bile acids prevent STAT3 phosphorylation induced by LIF inhibiting (altering/diminishing) the assembly of the LIFR/GP130 complex at cell membrane.

### LIFR antagonism exerts by DCA and LCA families limiting MKN45 cell proliferation and migration

To functionally characterize the effect of DCA and LCA families as LIFR antagonists, we conducted several *in vitro* assays using the poorly differentiated human GC cell line, MKN45. As previously demonstrated, MKN45 exhibits high levels LIF and LIFR expression^44^, making it an ideal candidate for our investigation.

First, we have explored whether DCA and its conjugated members, including TDCA, GDCA, and 3-oxoDCA, could modulate the proliferation of GC cells, mediated by LIF. MKN45 cells, cultured in serum-free medium, were exposed to 10 ng/mL LIF alone or in combination with increasing concentrations of DCA, TDCA, GDCA, and 3-oxoDCA (1-3-10-20 μM) for 8 hours. Cell vitality was assessed using the MTS assay as detailed in Material and Method extended section. Figure 3A shows that DCA did not reverse the LIF-induced proliferation, while TDCA exhibited a concentration-dependent reduction in cell vitality. GDCA and 3-oxoDCA reduced LIF-induced proliferation at 3 μM, but not at higher concentrations.

**Figure 3.**
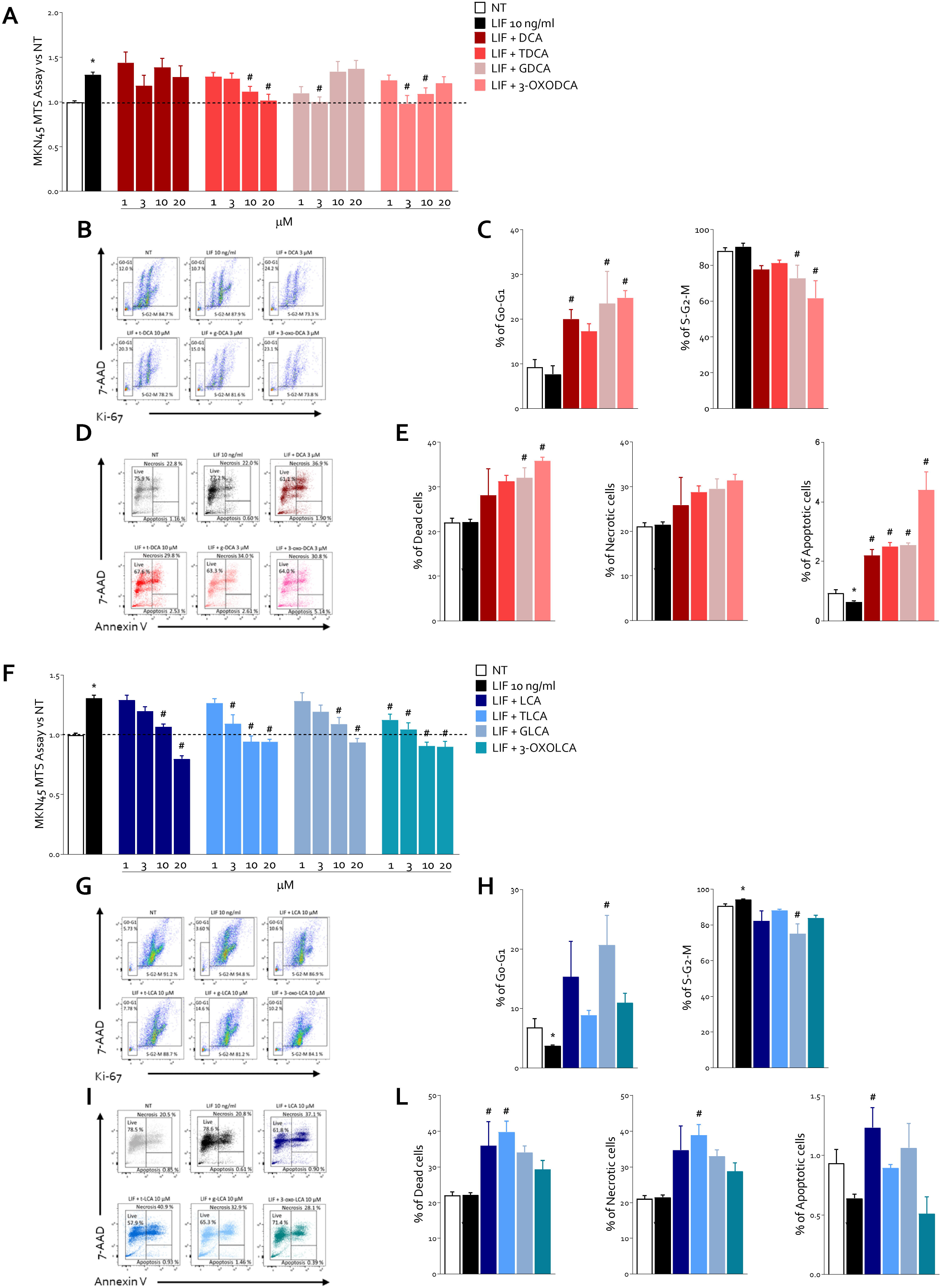
DCA and LCA family reversed cell proliferation rate induced by LIF on MKN45 cells. MNK45 cell lines was exposed to LIF alone or plus DCA members or LCA family members for 24 h or left untreated. Proliferation cell rate was determined using MTS assay: **A)** Concentration-response curve of DCA and its T, G, and 3-oxo derivatives (1, 3, 10, 20 µM). Cell cycle phase analysis was performed by Ki-67/7-AAD staining through IC-FCM. **B)** Representative IC-FCM shows cell cycle fraction in NT, LIF 10 ng/mL and LIF plus DCA (3 μM), TDCA (10 μM), GDCA (3 μM) and 3-oxoDCA (3 μM) groups. Frequencies of cells in the **C)** G0-G1 phase and S-G2-M phase. **D)** Representative IC-FCM shows Annexin V+ cells in each experimental group. Data shown are frequencies of **E)** Dead single cells, Necrotic single cells and Apoptotic single cells. **F)** MTS assay of LCA and its T, G, and 3-oxo derivatives (1, 3, 10, 20 µM). **G)** Representative IC-FCM of cell cycle fraction in NT, LIF 10 ng/mL and LIF plus LCA and its T,G and 3-oxo derivatives (10 μM). **H)** G0-G1 phase and S-G2-M phase cell rates. **I)** Representative IC-FCM shows Annexin V+ cells in each experimental group. **L)** Dead single cells, Necrotic single cells and Apoptotic single cells frequencies. (* represents statistical significance versus NT, and # versus LIF, *p < 0.05).

The effect of DCA and its T, G and 3-oxo derivatives on cell replication was also evaluated by Ki-67/7-AAD IC-FACS staining (Figure 3 B-C). Our findings demonstrated that challenging MKN45 with DCA and its derivatives modulates the cell cycle progression. More specifically, while LIF increased the transition into S-G2-M cell cycle phase, this effect was reversed by the exposure to 3 μM of DCA, GDCA and 3-oxoDCA, leading to increased frequencies of cells in G0-G1 resting phase, thereby blocking the transition to the replicative S-G2-M cell cycle phase, in a statistically significant manner (Figure 3 C).

Furthermore, we have shown that DCA, TDCA, GDCA and 3-oxoDCA modulate the apoptosis cell rates, as assessed by Annexin V/7-AAD staining (Figure 3 D).

Similarly, we have investigated the impact of LIFR inhibition mediated by LCA and its T, G and 3-oxo derivatives on the growth and proliferation of GC cells. For this purpose, MKN45 were cultured in a serum free medium, and exposed to 10 ng/mL LIF alone or in combination with increasing concentrations of above-mentioned natural bile acids (1, 3, 10, 20) for 8 h. As shown in Figure 3E, LCA reversed the LIF-induced proliferation at 10 μM concentration, while TLCA reduced cell vitality at the 3 μM concentration, GLCA blocked cells proliferation at 10 μM and 3-oxoLCA reversed LIF-induced growth even at the lowest concentration of 1 μM, as measured by the MTS assay. Only GLCA modulated the cell cycle progression blocking the shift from the G0-G1 resting phase to the replicative S-G2-M phase, as observed in the Ki-67/7-AAD IC-FACS staining results (Figure 3 G-H). LCA increased the frequencies of apoptotic cells and TLCA of improved the percentage of necrotic cells as determined by Annexin V/7-AAD staining (Figure 3 I-L).

To better characterize the molecular effect exerted by DCA and its derivatives as LIFR antagonists, we have evaluated the relative mRNA expression of several pro-oncogenic markers. As shown in Figure 4 A, whether the exposure of MKN45 to 10 ng/mL of LIF alone for 24 h increased the expression of the oncogene CMYC, the EMT marker Vimentin, and the anti-apoptotic BCL2, solely the treatment of LIF - exposed MKN45 with GDCA reduced the expression of CMYC and Vimentin. Similarly, the LIF-induced over-expression of BCL2 was reverted by the exposure to GDCA and 3-oxoDCA.

**Figure 4.**
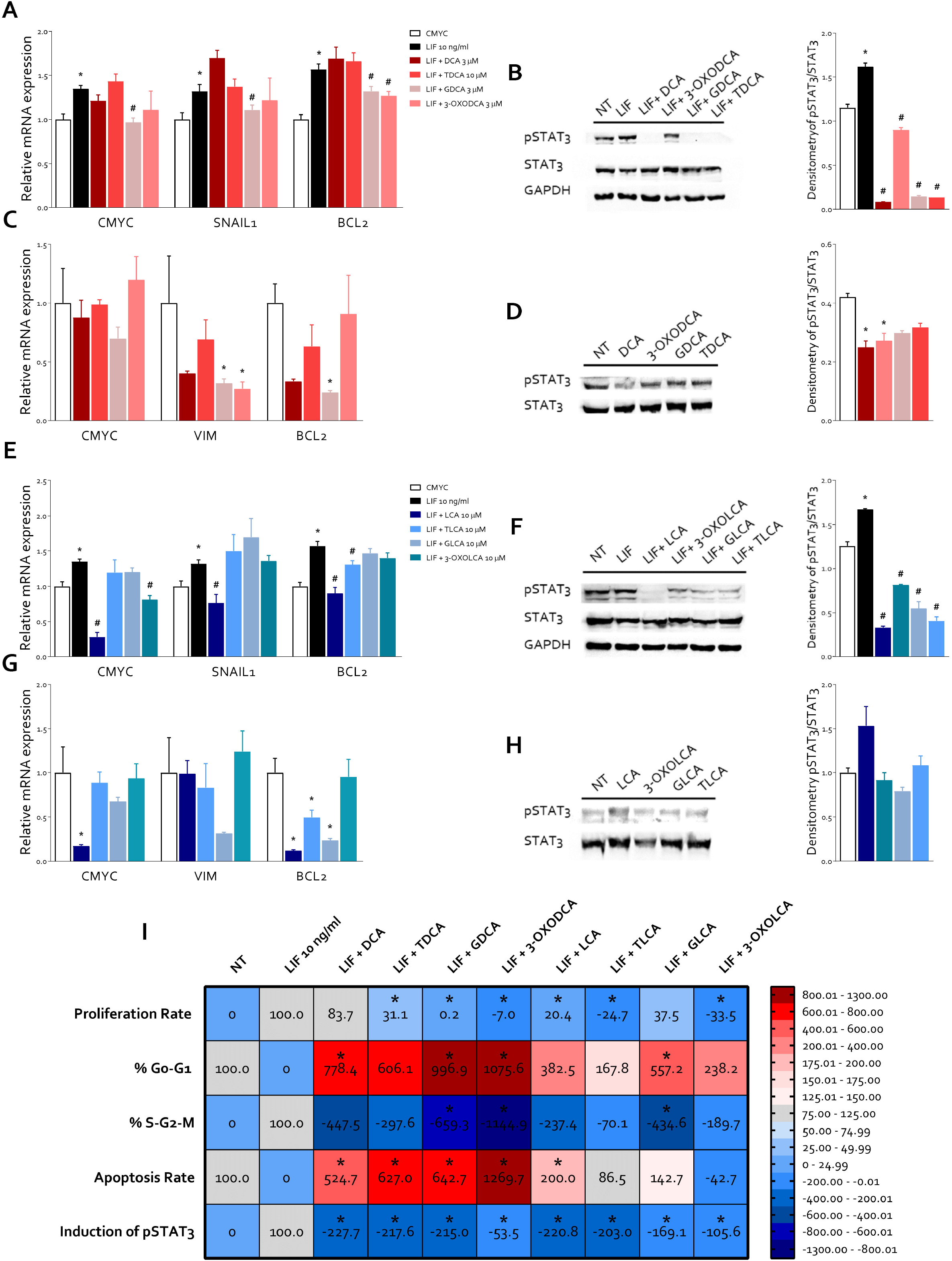
DCA and LCA family members reverse LIF-induced cancer features. In an experimental set, MKN45 cell line was exposed to LIF alone or plus DCA family members or LCA family members for 24 h or left untreated. In another experimental set, MKN45 cell line was exposed to DCA or LCA family members alone or left untreated. Relative mRNA expression of **A)** CMYC, SNAIL1 and BCL2 in MKN45 cells left untreated or exposed to LIF (10 ng/mL) alone or plus DCA and its T,G, and 3-oxo derivatives. Each value is normalized to GAPDH and is expressed relative to those of NT, which are arbitrarily set to 1**. B)** Representative Western blot analysis of phospho-STAT3 and STAT3 proteins (on left) and densitometric analysis demonstrating phospho-STAT3/STAT3 ratio (on right). **C)** Relative mRNA expression of CMYC, VIM and BCL2 in MKN45 cells left untreated or exposed to DCA and its T,G, and 3-oxo derivatives. Each value is normalized to GAPDH and is expressed relative to those of NT, which are arbitrarily set to 1**. D)** Representative Western blot analysis of phospho-STAT3 and STAT3 proteins (on left) and densitometric analysis demonstrating phospho-STAT3/STAT3 ratio (on right). Relative mRNA expression of **E)** CMYC, SNAIL1 and BCL2 in MKN45 cells left untreated or exposed to LIF (10 ng/mL) alone or plus LCA and its T,G, and 3-oxo derivatives. Each value is normalized to GAPDH and is expressed relative to those of NT, which are arbitrarily set to 1. **F)** Representative Western blot analysis of phospho-STAT3 and STAT3 proteins (on left) and densitometric analysis demonstrating phospho-STAT3/STAT3 ratio (on right). **G)** Relative mRNA expression of CMYC, VIM and BCL2 in MKN45 cells left untreated or exposed to LCA and its T,G, and 3-oxo derivatives. Each value is normalized to GAPDH and is expressed relative to those of NT, which are arbitrarily set to 1. **H)** Representative Western blot analysis of phospho-STAT3 and STAT3 proteins (on left) and densitometric analysis demonstrating phospho-STAT3/STAT3 ratio (on right). **I)** Heatmap of correlation summarizes cancer feature analyzed above.

Since LIF/LIFR modulates JAK and STAT3 phosphorylation ^57^, we have then investigated whether DCA and its derivatives could reverses the phosphorylation of STAT3, caused by LIF. Western blot assays, displayed in figure 4B, showed that all DCA family members reduced STAT3 phosphorylation in a statistically significant manner, confirming their role as suppressors of LIF/LIFR axis (Figure 4 B).

The role of secondary bile acids in cancer initiation and progression has traditionally been considered ambiguous. While several studies reported their oncogenic potential in gastrointestinal cancer development, more recent research has supported their robust anti-tumor effect ^58^. In this context, we investigated whether DCA and its derivatives might exert an oncogenic effect. We demonstrated that the exposure of MKN45 to DCAs did not promote their growth and proliferation. Specifically, bile acids had no effect on CMYC gene expression. GDCA and 3-oxoDCA downregulated the mRNA expression of Vimentin compared to untreated cells and the expression of BCL2 was reduced by GDCA (Figure 4 C). These findings indicated that bile acids did not activate the LIF/LIFR pathway and DCA and 3-oxoDCA reduced STAT3 phosphorylation (Figure 4 D).

Similar analyses were carried out to define the action of LCA family members as LIFR inhibitors. For this purpose, MKN45 were exposed to 10 ng/mL of LIF alone or in combination with LCA and its derivatives for 24 h. We found that LCA and 3-oxoLCA reduced the relative mRNA expression of CMYC. In addition, SNAIL-1 gene expression was reverted by LCA exposure, also LCA and 3-oxoLCA downregulated the mRNA expression of the anti-apoptotic BCL2 compared to LIF-challenged MKN45 (Figure 4 E).

LCA and its derivatives reduced STAT3 activation induced by LIF (Figure 4 F). Furthermore, the treatment of MKN45 with LCA and its derivatives alone did not promote a pro-oncogenic effect. As shown in Figure 4 G, LCA reduced the expression of CMYC compared to untreated cells and the relative mRNA expression of BCL2 was diminished by LCA, TLCA and GLCA. In addition, LCA and its derivatives had no effect on STAT3 activation pathway (Figure 4 H).

The results of these experiments were summarized in the heat map shown in Figure 4 I. The fifth column of the heat map shows that GDCA had the strongest effects on GC cell line proliferation, cell cycle regulation, apoptosis rate and STAT3 phosphorylation compared to the other bile acids tested.

### Bile acids content was increased in gastric neoplastic mucosa

Since these results demonstrate that secondary bile acids act as LIFR antagonists reversing the pro-oncogenic effects of LIF in GC cell lines, we have investigated the tissue content of various bile acids in paired surgical samples obtained from seven GC individuals and compared them with paired samples of macroscopically normal mucosa. These patients were selected from a larger cohort of GC patients (2013-2022) based on the availability of clinical and histological data, as well as paired tissue samples from non-neoplastic and primary neoplastic tissues.

The results of these studies demonstrated that GC samples harboured a significantly higher concentration of total bile acids compared to the non-neoplastic pairs while as much as 27 bile acids were identified (Figure 5 A). Among the primary bile acids, concentrations of GCA and GCDCA were significantly higher in GC samples than non-neoplastic pairs (Figure 5 B), while the percentage of HCA was decreased. In contrast, as shown in Figure 5 C, the percentage of secondary bile acids GDCA and 3-oxoDCA was significantly reduced in GC samples compared to non-cancer pairs (Figure 5 C). Together these data suggest that, while the total amount of bile acids increases in GC samples in comparison to non-cancer pair samples, the amount of GDCA and 3-oxoDCA, two LIFR antagonists, decreases, possibly indicating an inverse correlation among these bile acids species and LIFR expression. To clarify this question, we have then carried out a correlation analyses between the tissue contents of various bile acids and the tissue expression of LIFR (Log2), as derived from the transcriptome analysis described in a previous study (doi: 10.17632/7j7vm89d96.1) from which our cohort of seven GC patients was derived. As depicted in Figure 6 A-C, GDCA correlates negatively with LIFR expression in non-neoplastic mucosa, but this correlation was lost in neoplastic mucosa, due to the reduction of GDCA content in the cancer tissues (Figure 6 A-C). Additionally, 3-oxoDCA exhibited a linear correlation with LIFR expression in non-neoplastic mucosa but displayed an inverse correlation with neoplastic mucosa, similar to LCA (Figure 6A-C). These findings indicate that in non-cancer mucosa, but not in cancer pairs, a negative correlation occurs between GDCA concentrations and LIFR expression.Since these changes might contribute to a dysregulated LIF/LIFR pathway and oncogenesis in GC, we have then examined whether GCA, a bile acid whose concentrations are increased in GC samples, and GDCA, a bile acid whose concentrations are reduced in GC samples in comparison with non-cancer pairs, exerted a divergent effect on expression of pro-oncogenic markers of MKN45 cells challenged with LIF. As shown in Figure 5 E, while exposure of cells to LIF increased the expression of CMYC and SNAIL1, two oncogenetic markers, these effects were reversed by GDCA but not by GCA.

**Figure 5.**
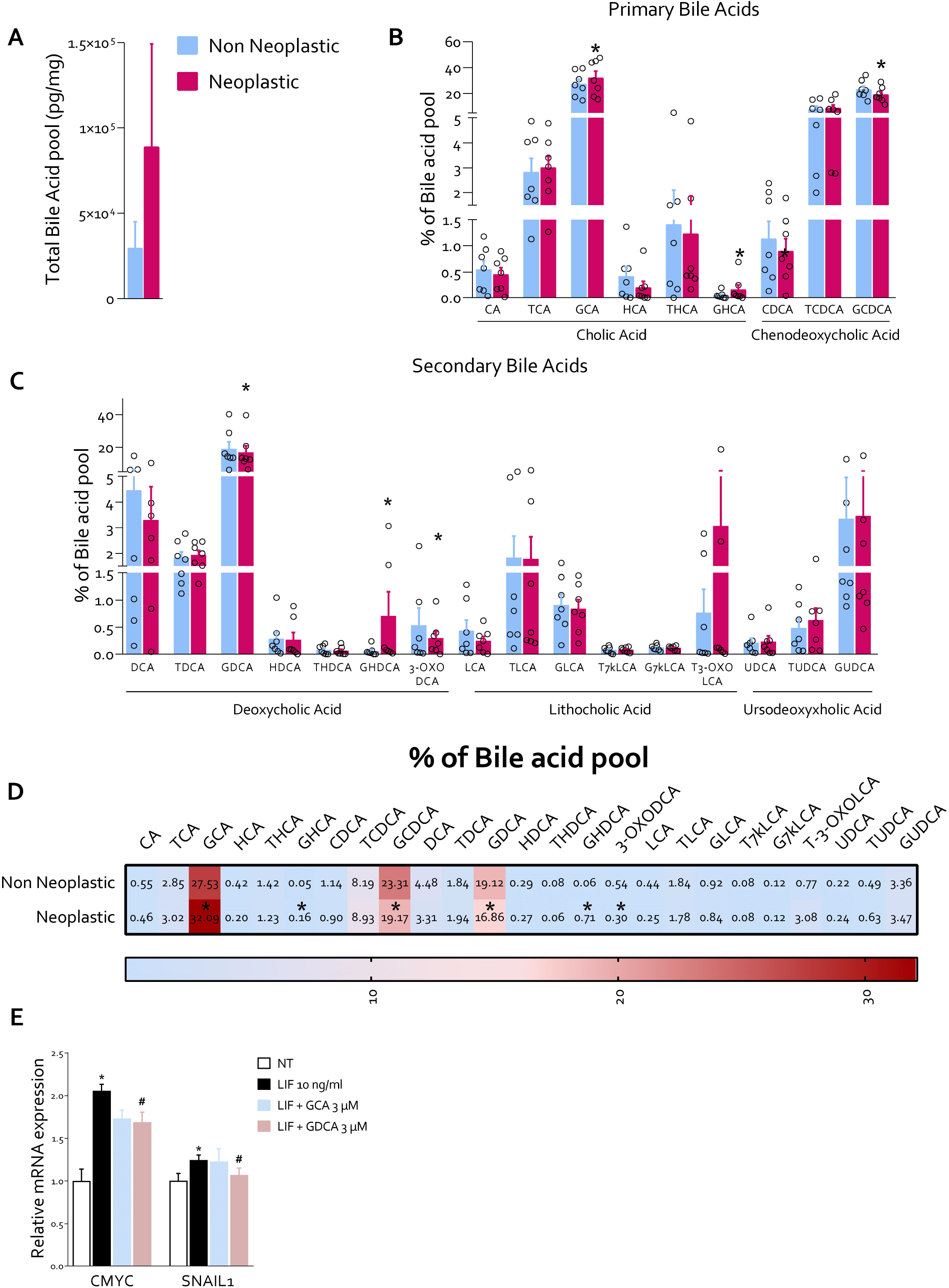
GDCA appears reduced in human gastric neoplastic mucosa compared to non-neoplastic mucosa. **A)** Total amount of bile acids pool (pg/mg) in non-neoplastic and in neoplastic mucosa. **B)** Percentage of primary bile acids pool in non-neoplastic and neoplastic mucosa. **C)** Percentage of secondary bile acids pool in non-neoplastic and neoplastic mucosa. **D)** Correlation Heatmap of percentages of bile acids pool between non-neoplastic and neoplastic mucosa. **E)** Relative mRNA expression of CMYC and SNAIL1 in MKN45 exposed to LIF (10 ng/mL) alone or plus GCA (3 μM) and GDCA (3 μM). Each value is normalized to GAPDH and is expressed relative to those of NT, which are arbitrarily set to 1**. (*p < 0.05).**

**Figure 6.**
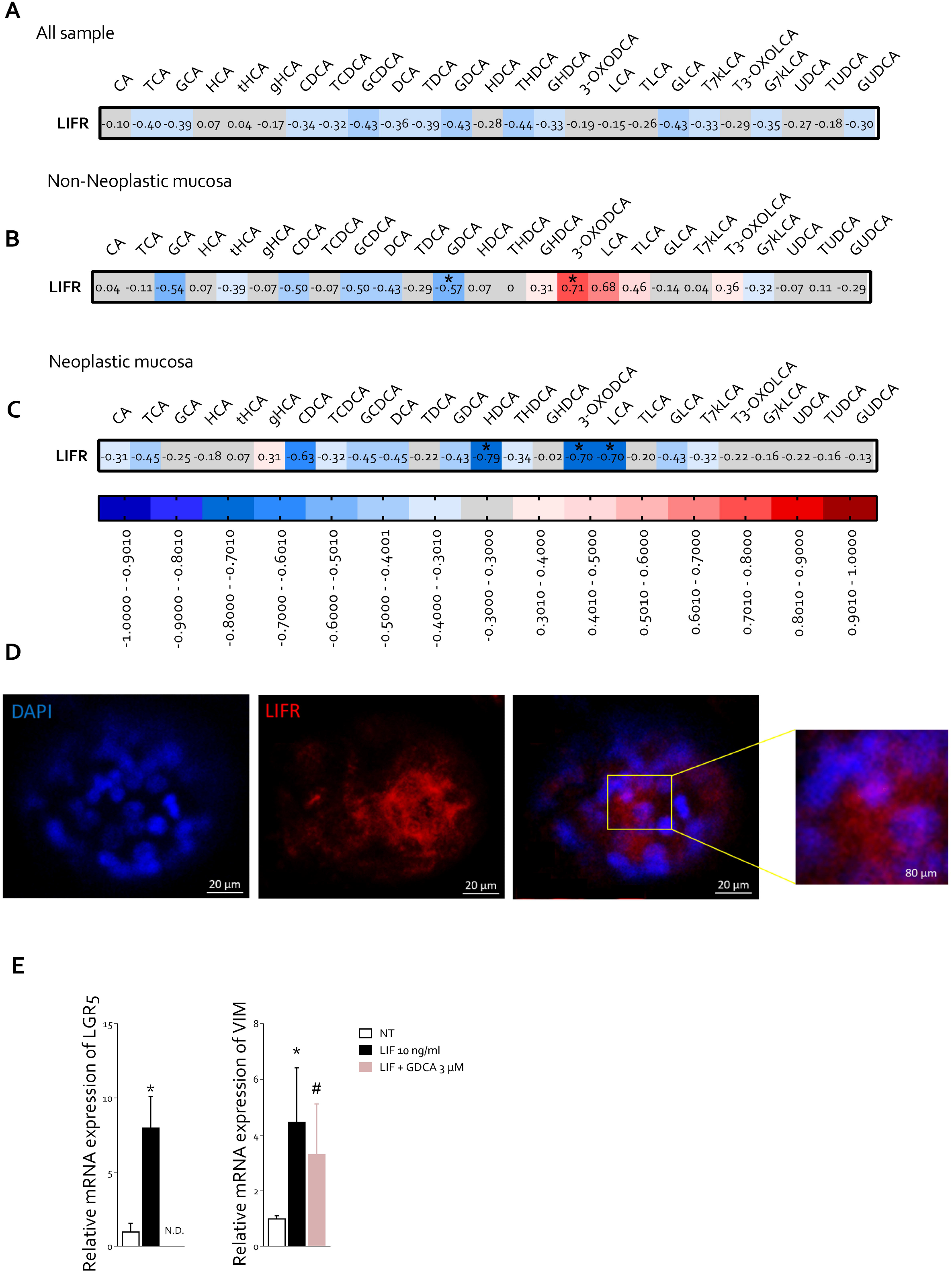
LIFR expression was inversely correlated with GDCA concentration in non-neoplastic mucosa but not in neoplastic mucosa. Heatmap of correlation between LIFR expression (Log2) and bile acid concentration (pg/mg) in **A)** All sample **B)** Non-neoplastic mucosa **C)** Neoplastic mucosa. **(*p < 0.05). GDCA inhibits the LIF-induced stemness properties of CSCs in hPDOs D)** Immunofluorescence staining of LIFR (red) in hPDOs (Magnification 20 μM and 80 μM) **E)** Relative mRNA expression of LGR5+ (on left) and VIM (on right).

To further validate these findings, patient-derived organoids (hPDOs) from GC patients were employed. Using these tissues, we were able to detect expression of LIFR on the cell membrane of hPDO (Figure 6D), as indicated by the intense red signal surrounding the DAPI-labelled nuclei. Challenging these hPDOs with 10 ng/mL LIF promoted a significant induction of the expression of the stem cell marker LGR5+ and Vimentin. These effects were reversed by GDCA (Figure 6E).

Similarly, while treating murine gastric organoids with LIF (10 ng/mL) alone or in combination with GDCA (3 µM) for 1 week promoted the growth of gastric organoids that appeared as enlarged masses formed by multiple layers of different cell types, these effects were reversed by GDCA (Supplementary Figure 2 B, C).

Collectively, these data demonstrate that bile acid species are differentially represented in cancer samples in comparison to their non-cancer pairs in GC and GDCA might exert a regulatory role on LIF/ LIFR signalling in GC.

### Secondary bile acids reverse the proliferative effect of LIF in pancreatic and colon cancer cell lines

Finally, we have investigated whether LIFR antagonism exerted by secondary bile acids in GC would extend to other cancers. Gastrointestinal cancer cell lines, MIA PaCa-2 (pancreas), HepG2 (liver) and Caco-2 (colon) cells, were used. As shown in Figure 7A and B, the expression of LIF/LIFR was detected in all cell lines by PCR analysis. While exposure to LIF promoted cells proliferation as assessed by MTS assay in all three cell lines, this pattern was reversed by TLCA, GLCA, 3-oxoLCA and TDCA (pancreas, liver and colon cancer). Also GDCA and 3-oxoDCA effectively reversed the LIF effects on liver and colon cancer cells, but were found less effective on MIAPaCa-2 cells (Figure 7C-E). Together these findings suggest that secondary bile acids function as LIFR antagonists and their tissue content correlates with LIFR expression in GC and their potential for LIFR antagonism might have relevance in regulating the oncogenic potential of this cytokine in pancreatic hepatic and colon cancers.

**Figure 7.**
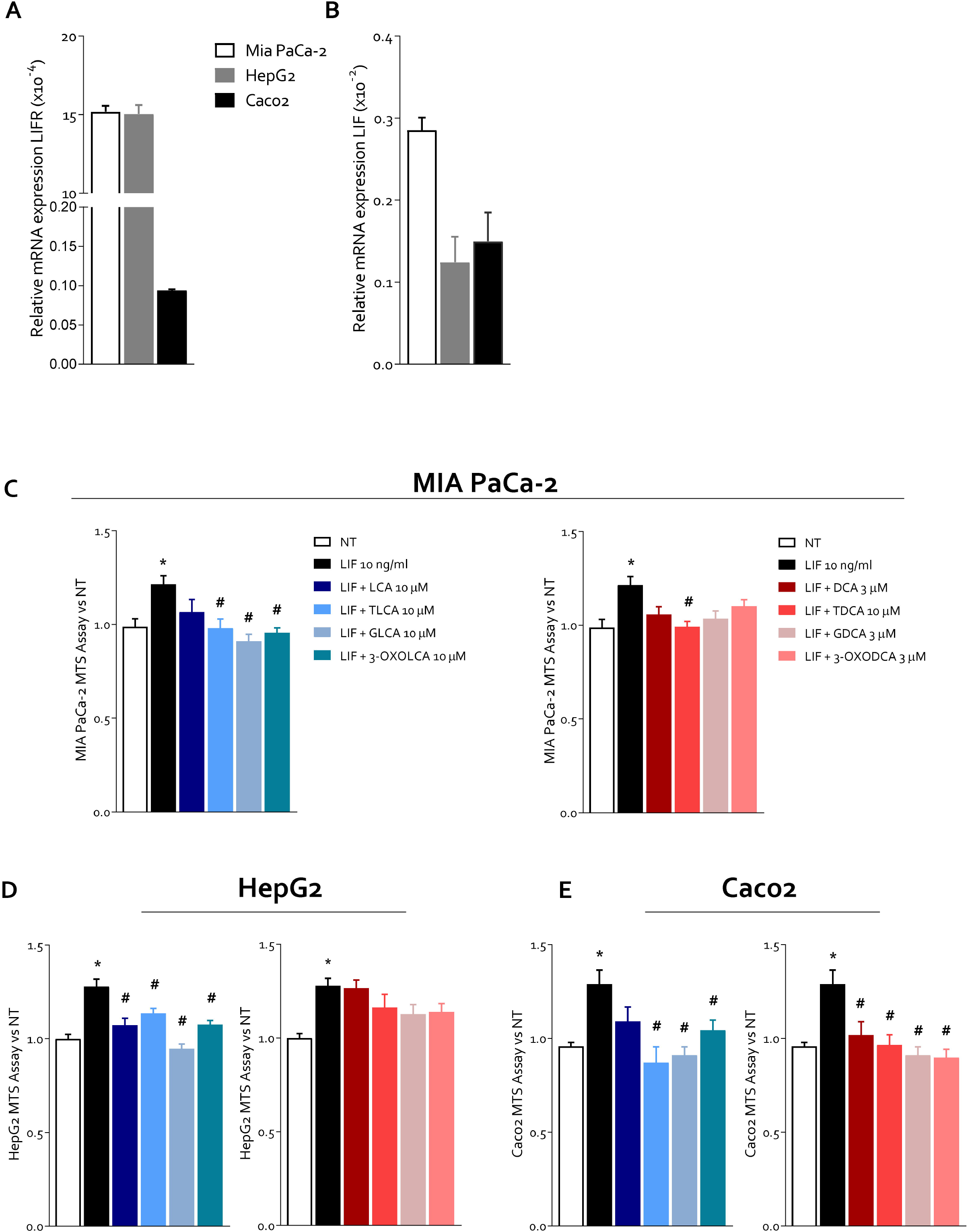
Secondary bile acids reversed proliferation rate LIF induced in gastrointestinal cell lines. Relative mRNA expression of **A)** LIFR and **B)** LIF in pancreatic MIA-PaCa2 cells (white), hepatic HepG2 cells (grey) and intestinal Caco2 cells (black). MTS assay on **C)** MIA PaCa-2 **D)** HepG2, **E)** Caco2 exposed to LIF alone or plus LCA and its T,G; and 3-oxo derivatives (shades of blue) and DCA and its derivatives (shades of red). (* represents statistical significance versus NT, and # versus LIF, *p < 0.05).

## Discussion

Here we report that secondary bile acids, DCA and LCA, the metabolic products of the intestinal microbiota, act as endogenous LIFR antagonists and that a decreased content of GDCA in GC associates with LIF/LIFR activation. These results expand the repertoire of signaling molecules generated by the intestinal microbiota and on their role in maintaining human health.

Previous studies have shown that bile acids function as endogenous ligands for cell membrane and nuclear receptors. Primary bile acids are the natural ligands of FXR^18,59,60^, the main bile acid sensor, while secondary bile acids act as non-exclusive ligands of PXR^34^, VDR^35^ and LXRs^37^ and RORγt^38^. For all these receptors, bile acids act as direct agonists, including RORγt, a nuclear receptor expressed in Th17 cells^61^. In the latter case, 3-oxoLCA and iso-alloLCA function as inverse agonists leading to RORγt-dependent inhibition of Th17 signaling at the intestinal microbiota/intestinal barrier interface^62^. Since intestinal inflammation associates with intestinal dysbiosis and reduced generation of RORγt agonists^63^, these studies established a role for microbial-derived bile acids in regulating host immune response by tuning the balance of Th17 and T regulatory cells^38,62^. In addition to nuclear receptors, secondary bile acids are the endogenous agonists of GPBAR1^20^, a membrane receptor that is also activated by primary bile acids though at significant higher concentrations^20^. Secondary bile acids might also activate the M2/M3 muscarinic receptors in certain cancers^64^, indicating that bile acids might function as endogenous ligands for both G-protein coupled and nuclear receptors ^65^.

We have now extended on this background by showing that bile acids function as LIFR antagonists. Our introductive screening of primary and secondary bile acids by Alfa Screen analysis demonstrated that LIFR antagonism waspreferentially restricted to secondary bile acids.A detailed analysis of concentrations response curve using human cells transfected whose construct has been made up by LIFR cloned upstream to a STAT3 reporter genes, confirmed that DCA and LCA and their T and G conjugates, along with their 3-oxo derivates, effectively prevent STAT3 phosphorylation induced by LIF. Inhibition of LIF/LIFR interaction and STAT3 phosphorylation occurs in the low micromolar concentrations, that is compatible with activation of GPBAR1 by secondary bile acids. Previous studies, by this laboratory and others, ^20,66,67^ have demonstrated that activation of GPBAR1 by LCA and DCA occurs at EC50 values of 0.9-2.0 µM, while activation of FXR requires significantly higher concentrations (≈10-20 µM). Together, these data establish that secondary bile acids function as LIFR antagonists in the same range of concentrations required for the activation of GPBAR1. As such, DCA and LCA and their derivatives should be considered *bona fide* as dual GPBAR1 agonists and LIFR antagonists in tissues co-expressing both receptors.

Through Molecular Dynamics simulations, we have shown that both LCAs and DCAs enter a pocket located in the Ig-like domain D3 of the extracellular portion of LIFR. This pocket is generated by L2 and L3 loops, which are part of the LIF binding domain of LIFR and it is predicted that its occupation would impact on the ability of LIF to bind to its receptor. These computational studies were confirmed by co-immunoprecipitation studies showing that GDCA, reversed the formation of LIFR/gp13o complex induced by LIF. Together these studies strongly support the notion that occupation of the pocket formed at L2 /L3 loops prevents activation of LIFR by LIF leading to the release of LFR/gp130 complex.

The functional characterization of DCAs and LCAs on gastrointestinal cancer cell lines and gastric organoids has further confirmed that these bile acid species counteract the pro-oncogenic effects of LIF in same order of magnitude shown by cell free and transactivation assays. Thus, while DCA *per se* did not fully reverse the LIF proliferative effects, TDCA, GDCA and 3-oxoDCA fully reversed the LIF-induced prooncogenic effects in various cancer cells lines^46^. Moreover, the exposure to DCA and its derivatives reversed the effect of LIF on cell cycle progression and cell survival in GC cell lines. Similarly, LCA and its derivatives reversed the pro-oncogenic effects of LIF as shown by measuring the cell cycle transition and the rate of apoptotic cells (Annexin V ^+^ cells). Additionally, some of the DCAs and LCAs blunted the expression of pro-oncogenic genes, including CMYC and BCL2 and robustly reduced STAT3 phosphorylation caused by LIF indicating that, among bile acids, GDCA antagonizes the pro-oncogenic activities of LIF in cancer cell lines.

Because these *in vitro* data suggested that intratumor content of bile acids might modulate the LIF/LIFR signalling at the epithelial cells/ECM interface, we have then characterized the bile acids content and composition in paired samples of non-neoplastic and neoplastic stomach from GC patients. The results of these studies demonstrated that the intra-tumour content of bile acids was significantly higher than in paired biopsies obtained from non-cancer areas. In comparison with their non-neoplastic counterparts, cancer biopsies were characterized by a significantly higher content of CA while tumour content of GCDCA, GDCA, GHDCA and 3-oxoDCA was reduced. Furthermore, analysis of LIFR expression in neoplastic and non-neoplastic samples demonstrated a negative correlation between the tissue content of GDCA and LIFR in non-neoplastic tissues, but this regulation was lost in the cancer samples due to the reduction of intra-tumour content of GDCA. The inverse correlation of LIFR expression and GDCA in normal and cancer tissues provides a strong support to translational relevance of this mechanism in GC.

In the neoplastic tissues, however, we have identified a negative correlation between the expression of LIFR and HDCA, LCA and 3-oxoDCA. Although the tissues expression gives no direct information on the status of LIF/LIFR signalling, it is noteworthy that the tissue content of LIFR antagonists (LCA and 3-oxoDCA) was negatively correlated with the expression of the receptor in GC samples. Because the tissue expression of LIF/LIFR associates with development of peritoneal metastasis^44^ and predicts worse prognosis in GC patients^68^, these human data highlight the therapeutic potential of DCAs and LCAs in regulating the LIF/LIFR system in clinical settings. Further confirming the translational relevance of our findings, we have shown that GDCA, the only bile acid that is reduced in GC samples, reversed the induction of the expression of GC stemness markers, LGR5, and the expression of EMT marker induced by LIF in patient-derived ^31,69,70^. The same regulatory effect of GDCA on LIF/LIFR pathway was confirmed in murine gastric organoids exposed to LIF. Finally, we have shown that LIFR antagonism by LCAs and DCAs is not restricted to GC but could be demonstrated in other gastrointestinal cancer cell lines, including pancreatic^46,71^, liver and colon cancer cells^5^, thereby establishing that secondary bile acids are LIFR antagonists and that these effects are maintained across various cancer types in enterohepatic tissues.

The reason why cancer tissues harbour higher bile acids contents in comparison to non-neoplastic tissues was not investigated in this study. Several mechanisms might be involved. Indeed, all GCs included in this study were of the intestinal subtype, according to Lauren’s classification^72,73^. This specific histologic subtype is thought to progress from gastric intestinal metaplasia frequently associated to the *H. Pylori* infection^70,74^. As, in contrast to gastric epithelial cells, the intestinal epithelial cells import bile acids^75^, this might explain the higher content of various bile acids in the cancer tissues, although other explanations, including a specific microbiota composition, should be considered ^76^.

In summary, through Alpha Screen assays, molecular modelling, and pharmacological characterization, we have demonstrated that LCAs and DCAs function as endogenous LIFR antagonists in the same range of concentrations required for activation of GPBAR1. Combining data of LIFR expression and analysis of intra-tumour bile concentrations, we have identified GDCA as a putative negative regulator of LIFR expression and activity in GC patients. These findings expand our understanding of the intricate interplay between bile acids, regulatory cytokines, and cancers in the gastrointestinal tract.

## Acknowledgments

Not applicable.

## Author Contribution

Conceptualization S.F., A.Z., B.C.; Experimental design C.D.G, M.B., S.M, A.L. E.M.; data collection, C.D.G, R.B., C.M., G.U. Ma.B.; Chemical synthesis V.S. and C.F., AlphaScreen and UPLC analysis, E.M., M.C.M., V.S; molecular docking studies, A.L., F.M., B.C.; data analysis, C.D.G., M.B. and S.M.; gastric biopsies supply, N.N., L.G. A.D.; writing original manuscript draft, C.D.G., E.M., A.L. and S.F.; editing manuscript C.D.G., E.M., M.C.M, V.S., A.L., B.C., A.Z. and S.F.

## Declaration of interest

The authors declare no conflict of interest, financial or otherwise.

## Funding

This work was partially supported by grant from the Italian MIUR/PRIN 2017 (2017FJZZRC) and MIUR ITALY PRIN 2022 PNRR P20227JB3W.

## Resource availability

### Lead contact

Further information and requests for resources and reagents should be directed to and be fulfilled by the Lead Contact Stefano Fiorucci. Institutional ad funding agency requirements for resource and reagent sharing will be followed.

### Materials availability

This study utilized data derived from transcriptome analysis and bile acids analysis of gastric cancer biopsies from our cohort of patients, in accordance with permits FI00001 (n. 2266/2014) and FI0003 (n. 36348/2020). This resource is available upon request to the lead contact as indicated above. Institutional and funding agency requirements for resource and reagent sharing will be followed.

### Data and code availability

- All data reported in this paper will be shared by the lead contact by request.
- This paper does not report original code
- Any additional information required to reanalyze the data reported in this paper is available from the lead contact upon request

## Supplementary Figure legends

**Figure S1.** Gastric organoids were established from healty C57BL6/J mice. Data shown are: **A)** H&E staining of gastric organoid (on left). IF analysis of E-CADH (green) and LIFR (red) basal expression (on right). Gastric organoids were exposed to LIF (10 ng/ml) alone or in combination with GDCA (3 µM) for 1 week. **B)** Representative photos of 3D cultures of the three experimental groups. **C)** Number of single cells derived from 3D culture dissociation. (* represents statistical significance versus NT, and # versus LIF, *p < 0.05).

**Figure S2. A)** Ligand Root Means Square Deviation (L-RMSD) plot after 150 ns of MD simulation. **B)** Clusters distribution after 150 ns of MD simulations.

**Figure S3.** 3D and 2D views of the most representative clusters after 150 ns of MD simulation of the complex of hLIFR with DCA, GDCA, 3-oxoDCA, TDCA, TLCA and 3-oxoLCA. The binding site is defined by three loops, namely L1 (255-VSASSG-260), L2 (303-NPGRVTALVGPRAT-316), and L3 (332-KRAEAPTNES-341). Ligands and the main residues involved in the binding mode are labelled and visualized in the stick while the hydrophobic (HYD), H-Bond Acceptor and Donor (HBA/HBD) and the Negative Ionizable (NI) pharmacophore features are coloured as summarized in the caption. The H-Bond are highlighted in black dashed lines.

## Notes

https://data.mendeley.com/drafts/7j7vm89d96

### Competing Interest Statement

The authors have declared no competing interest.

