## Supplemental Table 1 and 2; Supplemental Figure 1, 2 and 3 for "Bile Acids Serve As Endogenous Antagonists Of The Leukemia Inhibitory Factor (LIF) Receptor in Oncogenesis": Supplementary Tables.docx

**TABLE S1.** QPLD and IFD docking results.

| **Compound** | **QPLD** | **IFD** | |
| --- | --- | --- | --- |
|  | *G-Score** | *G-Score** | *IFD-Score** |
| **DCA** | -4.685 | -9.426 | -10636.22 |
| **GDCA** | -4.129 | -9.745 | -10711.92 |
| **TDCA** | -4.389 | -7.956 | -10655.43 |
| **3-OXODCA** | -4.344 | -7.531 | -10610.95 |
| **3-OXOLCA** | -4.480 | -7.130 | -10602.22 |
| **TLCA** | -4.453 | -10.311 | -10726.18 |

* Value expressed in kcal/mol

**TABLE S2.** Cluster analysis and MM/GBSA ΔG value energy estimation of hLIFR-DCA (A), -GDCA(B), -TLCA (C), -3-OXODCA (D), -3-OXOLCA (E) and -TLCA (F) after 150 ns of MD simulation.

**A**

| **Cluster** | **% pop** | **AvgDist**^a^ | **Stdev** | **AvgCDist** | **MMGBSA**^b^ **(ΔG)**^c^ |
| --- | --- | --- | --- | --- | --- |
| 0 | 70 | 0.710 | 0.115 | 1.073 | -32.10 (±4.3) |
| 1 | 25 | 0.731 | 0.125 | 1.027 | -34.16 (±1.8) |
| 2 | 0.3 | 0.721 | 0.124 | 1.150 | - |
| 3 | >1 | 0.679 | 0.091 | 1.127 | - |
| 4 | >1 | 0.656 | 0.106 | 1.045 | - |

**B**

| **Cluster** | **% pop** | **AvgDist**^a^ | **Stdev** | **AvgCDist** | **MMGBSA**^b^ **(ΔG)**^c^ |
| --- | --- | --- | --- | --- | --- |
| 0 | 42 | 0.973 | 0.218 | 1.777 | -23.78 (±1.7) |
| 1 | 42 | 1.010 | 0.236 | 1.710 | -30.18 (±2.3) |
| 2 | 10 | 0.889 | 0.179 | 1.541 | -23.70 (±2.3) |
| 3 | >1 | 1.059 | 0.246 | 1.769 | - |
| 4 | >1 | 0.722 | 0 | 1.850 | - |

**C**

| **Cluster** | **% pop** | **AvgDist**^a^ | **Stdev** | **AvgCDist** | **MMGBSA**^b^ **(ΔG)**^c^ |
| --- | --- | --- | --- | --- | --- |
| 0 | 61 | 1.264 | 0.282 | 1.819 | -27.10 (±3.8) |
| 1 | 32 | 1.213 | 0.278 | 1.957 | -25.70 (±3.3) |
| 2 | >1 | 1.053 | 0.218 | 2.227 | - |
| 3 | >1 | 0.815 | 0 | 2.285 | - |
| 4 | >1 | 0 | 0 | 2.277 | - |

**D**

| **Cluster** | **% pop** | **AvgDist**^a^ | **Stdev** | **AvgCDist** | **MMGBSA**^b^ **(ΔG)**^c^ |
| --- | --- | --- | --- | --- | --- |
| 0 | 84 | 0.704 | 0.114 | 1.060 | -22.42 (±1.1) |
| 1 | 13 | 0.729 | 0.122 | 1.437 | -22.12 (±1.5) |
| 2 | >1 | 0.682 | 0.112 | 1.101 | - |
| 3 | >1 | 0.660 | 0.102 | 1.168 | - |
| 4 | >1 | 0.681 | 0.136 | 1.327 | - |

**E**

| **Cluster** | **% pop** | **AvgDist**^a^ | **Stdev** | **AvgCDist** | **MMGBSA**^b^ **(ΔG)**^c^ |
| --- | --- | --- | --- | --- | --- |
| 0 | 82 | 0.695 | 0.110 | 1.171 | -28.47 (±1.3) |
| 1 | >1 | 0.697 | 0.124 | 1.185 | - |
| 2 | >1 | 0.698 | 0.117 | 1.262 | - |
| 3 | >1 | 0.727 | 0.132 | 1.651 | - |
| 4 | >1 | 0.785 | 0.013 | 1.224 | - |

**F**

| **Cluster** | **% pop** | **AvgDist**^a^ | **Stdev** | **AvgCDist** | **MMGBSA**^b^ **(ΔG)**^c^ |
| --- | --- | --- | --- | --- | --- |
| 0 | 65 | 1.314 | 0.311 | 2.123 | -33.45 (±2.1) |
| 1 | 21 | 1.130 | 0.254 | 1.938 | -30.72 (±3.1) |
| 2 | >0 | 1.176 | 0.271 | 2.056 | - |
| 3 | >0 | 1.013 | 0.216 | 2.213 | - |
| 4 | >0 | 0.856 | 0.178 | 2.539 | - |

^a^AvgDst represents the maximum RMSD in Å from the other member of the cluster;  ^b^Calculated with CPPTRAJ module; ^c^ΔG expressed in *kcal/mol.*
