## Supplementary figures and images for "Bile Acids Serve As Endogenous Antagonists Of The Leukemia Inhibitory Factor (LIF) Receptor in Oncogenesis"

### Figure S1.tif

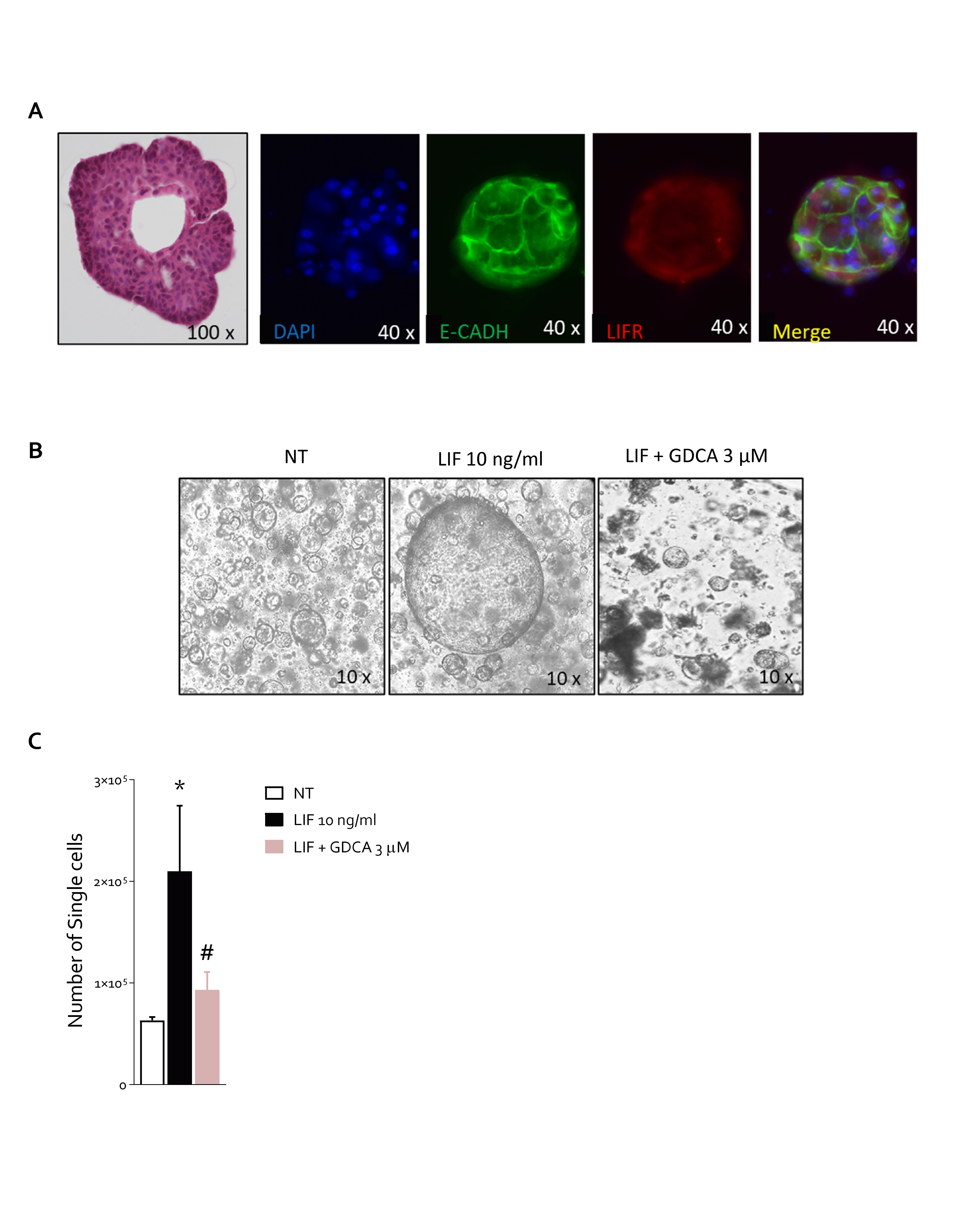

### Figure S2.tif

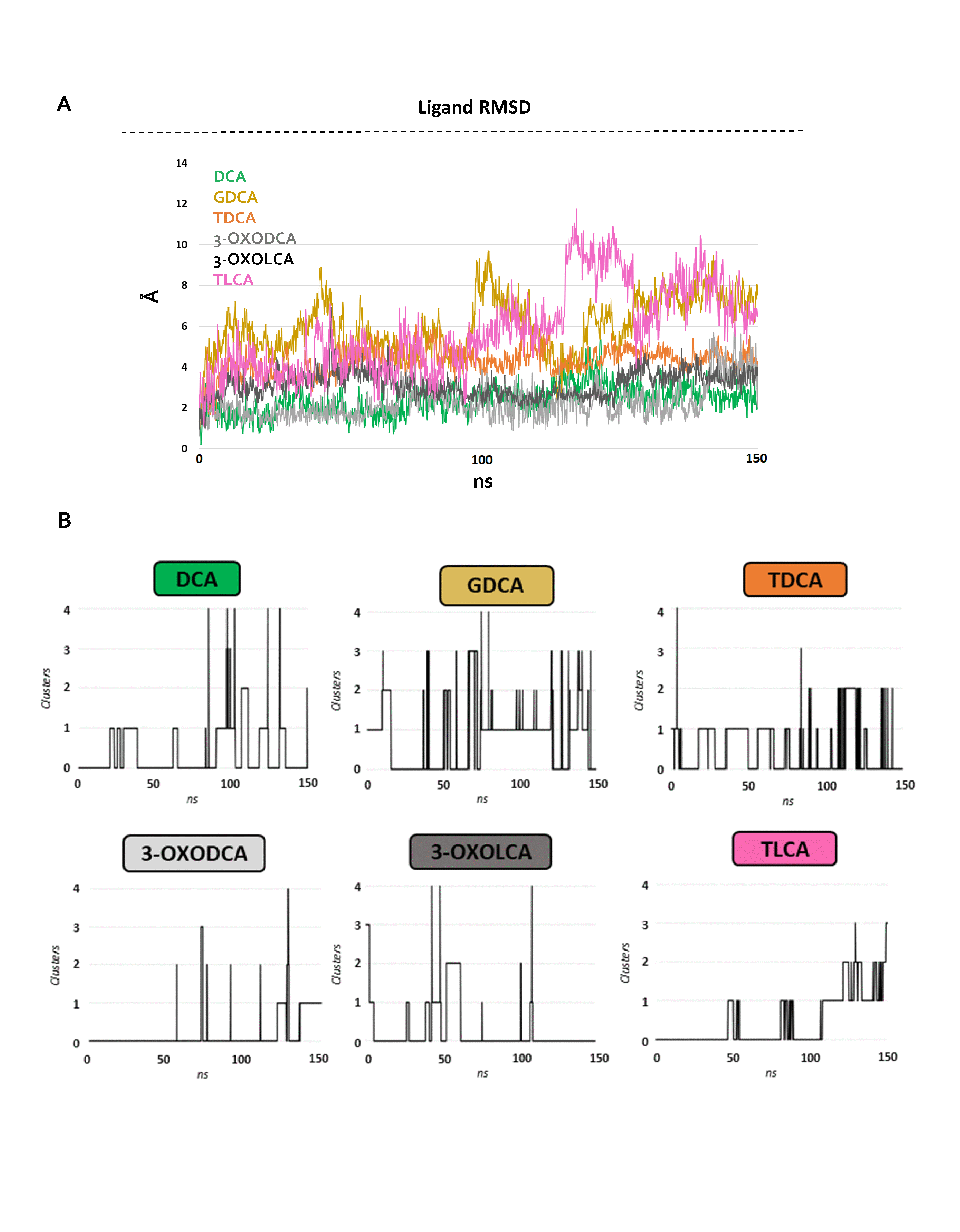

### Figure S3.tif

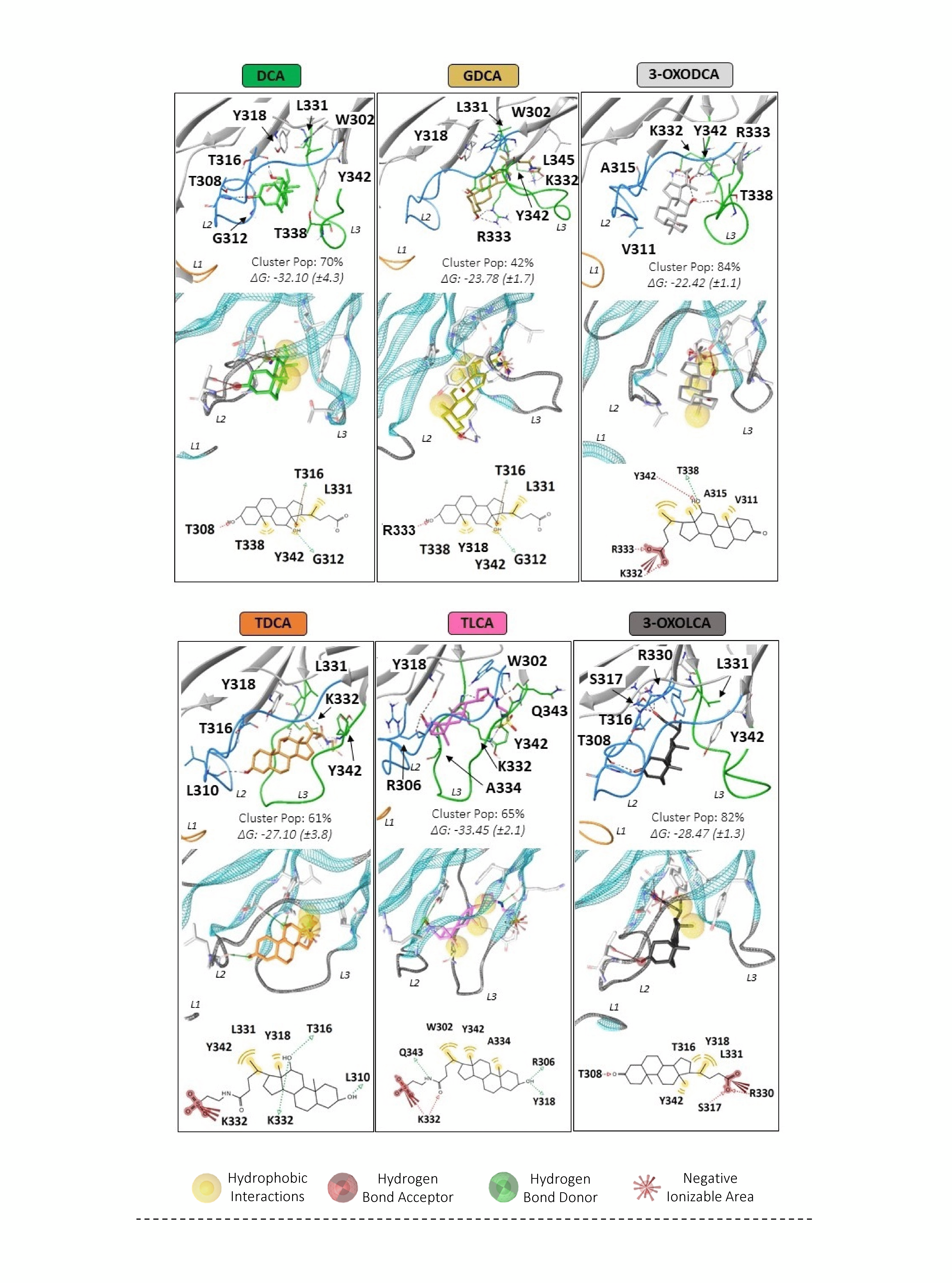
